# A combined lifestyle intervention induces a sensitization of the blood transcriptomic response to a nutrient challenge

**DOI:** 10.1101/2021.05.18.444591

**Authors:** Thies Gehrmann, Marian Beekman, Joris Deelen, Linda Partridge, Ondine van de Rest, Leon Mei, Yotam Raz, Lisette de Groot, Ruud van der Breggen, Marcel J. T. Reinders, Erik B. van den Akker, P. Eline Slagboom

## Abstract

The global population is growing older. As age is a primary risk factor of (multi)morbidity, there is a need for novel indicators to predict, track, treat and prevent the development of disease. Lifestyle interventions have shown promising results in improving the health of participants and reducing the risk for disease, but in the elderly population, such interventions often show less reliable or subtle effects on health outcomes. This is further complicated by a poor understanding of the homeodynamics and the molecular effects of lifestyle interventions, by which their effects of a lifestyle intervention remain obscured. In the Growing Old Together (GOTO) study, we examined the responses of 164 healthy, elderly men and women to a 13-week combined physical and dietary lifestyle intervention. In addition to collecting blood samples at a fasted state, we sampled blood also 30 minutes following a standardized meal. This allows us to investigate an intervention response not only in the traditional fasted state, but also in the blood metabolic and cellular responses to a nutrient challenge. We investigated the transcriptomic and metabolomic responses to this nutrient challenge, how these responses relate to each other, and how this response is affected by the lifestyle intervention.

We find that the intervention has very little effect on the fasted blood transcriptome, but that the nutrient challenge induces a large translational inhibition, and an innate immune activation, which together comprise a cellular stress response that is stimulated by the intervention. A sex-specific analysis reveals that although the same set of genes respond in the same direction in both males and females, the magnitude of these effects differ, and are modulated differently by the intervention. On the other hand, the metabolomic response to the nutrient challenge is largely unaffected by the intervention, and the correlation between the metabolomic nutrient response and transcriptomic modules indicates that the change in transcriptomic response to the nutrient challenge is independent from a change in cellular metabolomic environment.

This work constitutes a glance at the acute transcriptomic stress response to nutrient intake in blood, and how a lifestyle intervention affects this response in healthy elderly, and may lead to the development of novel biomarkers to capture the phenotypic flexibility of health.

## Introduction

The global population is growing older (WHO, 2015) and as age is a primary risk factor for many diseases (Niccoli and Partridge, 2012) society is obligated to adjust to this demographic shift. This adjustment may take the form of psychosocial or medical interventions, structural social changes that together remove the barriers that cause or exacerbate physical and mental disability, and the prevention or delay of morbidity and/or progression to multi-morbidity (WHO, 2015). Although ageing is a very personal process driven by extrinsic and intrinsic changes that differ from one individual to the next, it has become clear that healthy ageing can generally be stimulated by healthy lifestyles. Lifestyle interventions are known to reduce the risk of multiple conditions such as cardiovascular disease, diabetes, cancer, obesity and hypertension, and it is assumed that they will contribute to life- and healthspan (Fontana et al., 2010; Longo et al., 2015).

Lifestyle interventions that stimulate health include quitting smoking, reducing the consumption of alcohol and processed food, reduced nutrient intake or supplementation (Backx et al., 2016; Kraus et al., 2019; Layman et al., 2005), adopting physical and mental exercise (Pittas et al., 2006; Tieland et al., 2012), or combinations thereof (Josse et al., 2011; Layman et al., 2005; van de Rest et al., 2016; Schutte et al., 2016). While interventions have shown positive health effects, these are typically examined in men, the sick, the young, or those at very high risk, and often only in small studies. In contrast, for the elderly who are at most risk of age-related diseases, such interventions have less reliable or very subtle effects on health outcomes(Backx et al., 2016; Tieland et al., 2012, 2017). Additionally, the molecular basis of the individual response to interventions is still poorly understood. For instance, dietary or physical exercise interventions have shown changes in the metabolome (van Dijk et al., 2012; Pellis et al., 2012; Zeevi et al., 2015) but are frequently unable to identify changes in the transcriptome (Van Bussel et al., 2017; Fazelzadeh et al., 2018; ten Haaf et al., 2018). Also, the time-scale needed to investigate responses to interventions remains largely unknown. For instance, the effect of lifestyle interventions is increasingly investigated using dietary stress tests monitoring the response to overload lipid, protein or glucose administration. These metabolic and caloric challenge tests are regarded upon as novel potential biomarkers of health (van Ommen et al., 2014; Stroeve et al., 2015) recording the phenotypic flexibility of individuals within minutes to hours of the test triggering acute and complex stress responses (energy fluxes, oxidative and inflammatory stress, apoptosis etc.). Hence, there is a need to monitor the response not only by traditional health markers but also at the molecular level (Van Bussel et al., 2017; Catoire et al., 2012).

In the Growing Old Together (GOTO) study, we previously examined the response of 164 healthy, elderly men and women to a 13-week combined physical activity and dietary intake reduction lifestyle intervention (van de Rest et al., 2016; Schutte et al., 2016) and found that several indicators of health, including BMI, waist circumference and systolic blood pressure improved as a result. In addition we demonstrated a beneficial effect on the fasted serum metabolome exemplified by a decrease in glucose, lipids, branched chain and aromatic amino acids, inflammatory markers such as α-acid glycoprotein, largely independent of the weight change (van de Rest et al., 2016). Here, we complement these studies by investigating to what extent the blood transcriptome and serum metabolome response to a standardized nutrient intake (containing 35% fat, 49% carbohydrates, 16% protein) after an overnight fast may provide information on the phenotypic flexibility in older people. We examined to what extent a lifestyle intervention changed this response and how this response can be considered the substantiation of health effects due to the intervention. We find that the intervention has little effect on the unchallenged (fasted) transcriptome. In contrast, the transcriptomic response to the nutrient challenge provides a glimpse at an acute stress response in blood, that the intervention modulates the degree of this response, and does so in a sex-dependent manner. Moreover, we demonstrate that the metabolomic response to the nutrient challenge is largely not affected by the intervention, and that the transcriptomic effect of the intervention on the nutrient challenge constitutes a different cellular response to the same metabolomic environment.

## Results

### A 13-week combined lifestyle intervention improves the health of its participants

For 85 of the 164 GOTO (Materials and methods) participants, RNA sequencing data was collected from whole blood sampled before and after the 13-week combined dietary and physical lifestyle intervention (Materials and Methods). Sampling was conducted after an overnight fast (Figure 1A, timepoints 1 and 3) and again 30 minutes following administration of a standardized liquid meal (timepoints 2 and 4).

**Figure 1:**
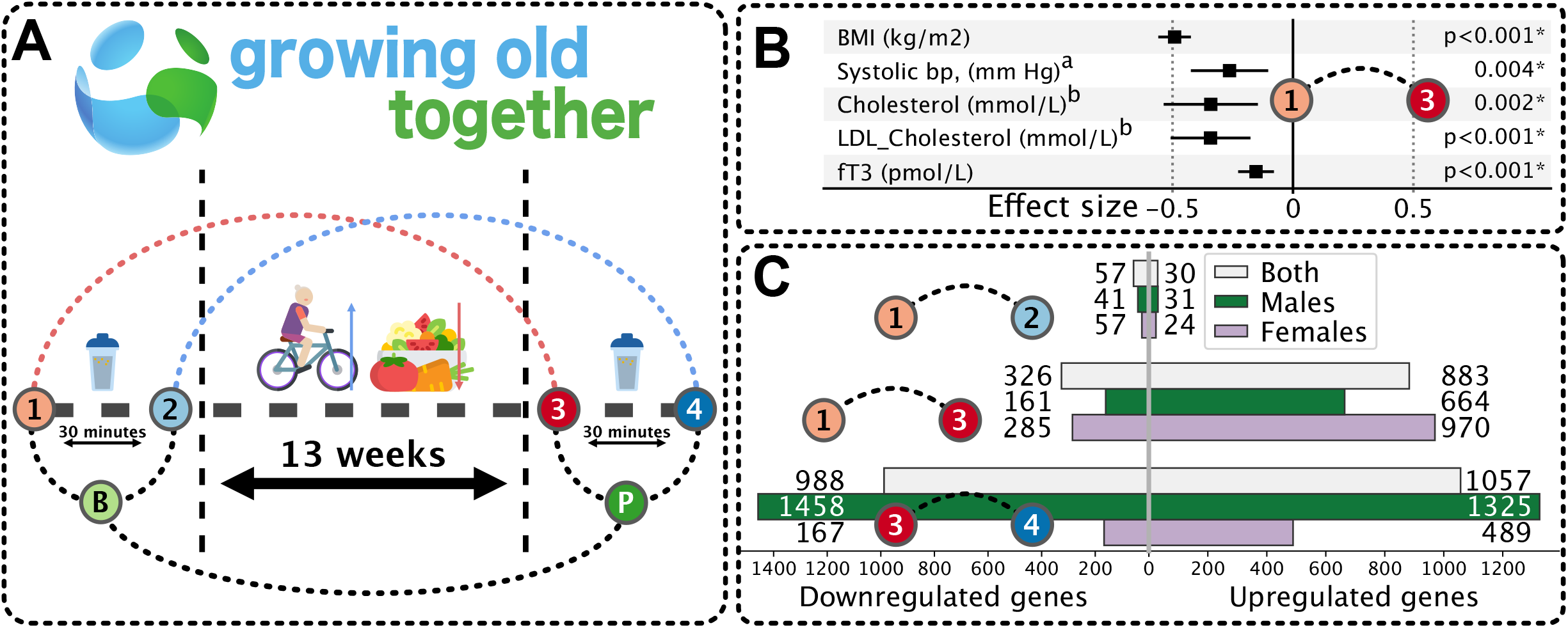
A. Outline of the GOTO study timepoints. Participants have blood sampled at four timepoints. Preceding the intervention in a fasted state (1), and 30 minutes following a standardized meal (2), and after the 13 week intervention in a fasted state (3), and 30 minutes following a standardized meal.
B. Health markers significantly improved as a result of the intervention in both males and females. The error bar indicates the 95% confidence interval. ^a^Individuals on antihypertensive medication were excluded from this analysis. ^b^Individuals on lipid-lowering medication were excluded from this analysis.
C. Boxplot of differential expression counts. Left: downregulated, Right: Upregulated. Top: Intervention effect in fasted tissues, for all samples (gray bars), and males (green) and females (purple) separately. Middle: Nutrient challenge effect at baseline Bottom. Nutrient challenge effect after the intervention

In previous studies, we demonstrated that health indicators such as parameters of body composition, physiological function, diagnostic serum parameters and disease related metabolites were positively affected by the combined lifestyle intervention. The intervention effects in these 85 individuals were comparable to those found in the complete study (van de Rest et al., 2016), finding that many of the most relevant health indicators for both men and women, including BMI, systolic blood pressure, cholesterol, HDL cholesterol and fT3 improved throughout the intervention (Figure 1B, SI Table 1). Only insulin showed a non-significant effect, though it was also the weakest detected effect when investigated in the whole study.

### Nutrient challenge raises a strong transcriptomic response and is enhanced by the intervention

To examine the effects of the intervention on the blood transcriptome, we performed a differential expression test, contrasting the fasted samples before and after the intervention. on both the fasted and postprandial samples (Materials and Methods, SI Table 2, SI Figure 1). This revealed that there were very few differentially expressed genes in the fasted or postprandial samples when baseline was compared to the intervention. In total, 87 genes were differentially expressed in the fasted samples (timepoints 3 vs 1), but no functional terms were enriched in this gene set.

While the intervention appeared to have a relatively small effect on the blood transcriptome, we observed large effects for each of the nutrient challenges. The nutrient challenge affected the postprandial expression of a large number of genes (Figure 1C), with 1209 differentially expressed at baseline (timepoints 2 vs. 1, Figure 1C), and 2045 following the intervention (timepoints 4 vs. 3, Figure 1C). We mark two observations: First, the set of postprandially downregulated genes after the intervention is larger than that at baseline (326 at baseline, 988 after, SI Table 2) and second, the set of genes postprandially upregulated after the intervention is largely the same as at baseline (883 before, 1057 after, overlapping with 617 genes,).

### Nutrient challenge affects a common set of genes in both sexes, with different magnitudes

As our study consisted of both men and women (44 and 41, respectively), we could investigate the male and female responses to the nutrient challenge separately. These analyses are rarely performed in challenge or intervention studies. In the sex-stratified analysis of the intervention effect in the fasted samples, the effects were still rather small in both males and females (72 and 50 genes, respectively).

In the sex-stratified nutrient challenge response, we observed remarkably different numbers in males and females both at baseline and after the intervention (Figure 1C, SI Table 2, SI Figure 1). Females exhibited more differentially expressed genes at baseline (1255, with 970 up-, and 285 down-regulated) than after the intervention (656 total, with 489 up−, and 167 down-regulated), males had fewer differentially expressed genes at baseline (825 total, with 664 up- and 161 down-regulated), and more differentially expressed after the intervention (2783 total, with 1325 up- and 1458 down-regulated after the intervention).

Generally speaking, the same gene is affected in the same direction by the nutrient challenge in both males and females, merely the magnitude is different (Figures 2A-B, SI Figure 2). Our observation indicates that the intervention affects the nutrient challenge response differently in males and females; in males the effect becomes more pronounced, whereas in females it becomes smaller. We conclude that we can define a set of genes that are downregulated or upregulated in response to a nutrient challenge in both males and females, both before and after the intervention. This set of genes consisted of 1904 upregulated, and 1721 downregulated genes.

**Figure 2:**
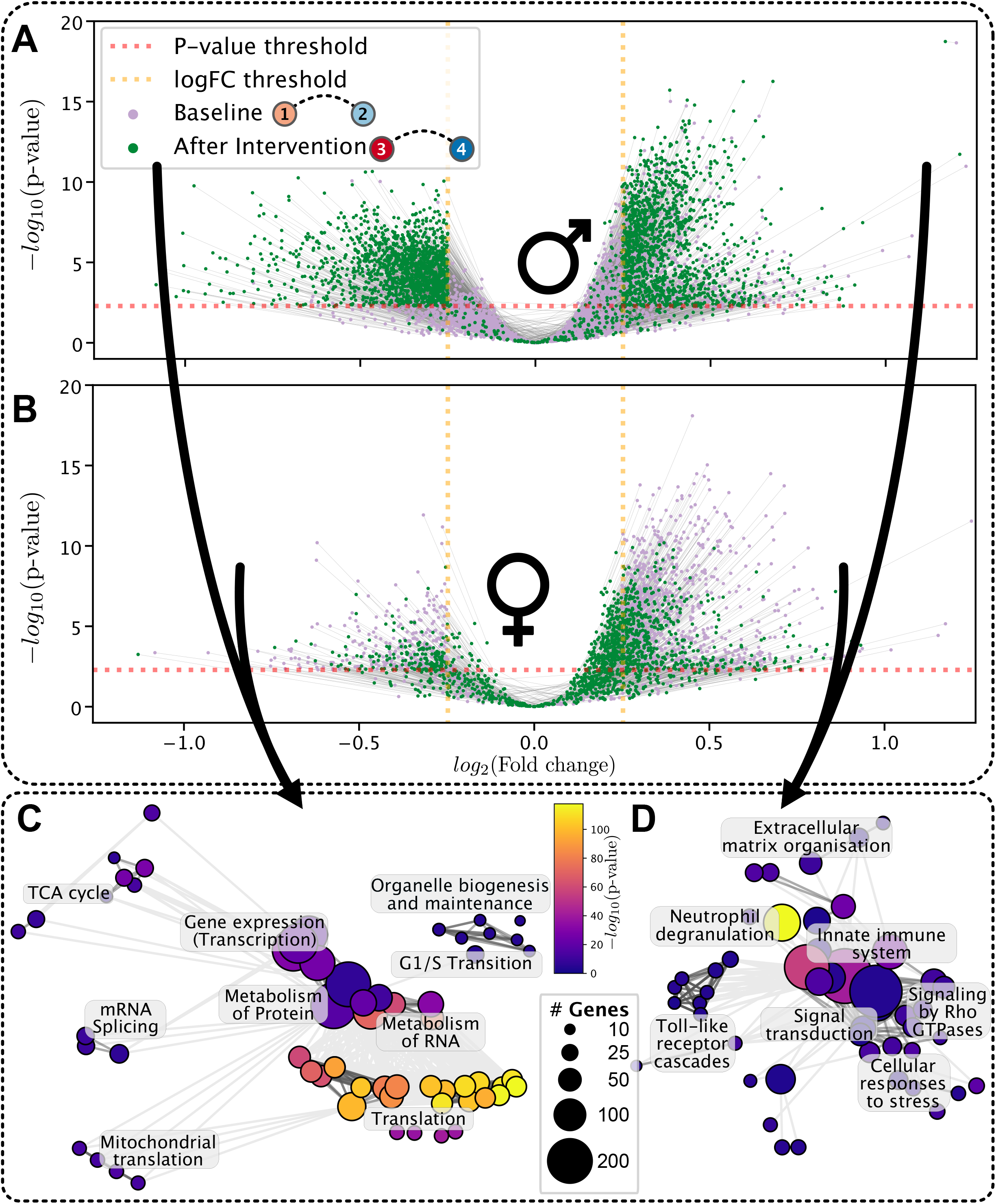
A. A paired-volcano-plot with results for males only. Each pair of points, connected by a line, represent the same gene at two conditions, the postprandial response, in purple at baseline, and in green following the intervention. Only genes which were significant in either of the two conditions are shown.
B. A paired volcano plot, as in 2A, but showing results for females only.
C. Functional enrichments in the consistently downregulated genes of the postprandial response. Each point refers to a Reactome annotation, and the distance between annotations is determined as the inverse of their gene overlap. Clusters are identified with a hierarchical clustering. The color of the point represents the −log10 FDR adjusted p-value of the enrichment, and the size represents the number of differentially expressed genes in that term.
D. Functional enrichment graph in the upregulated genes of the postprandial response.

### Nutrient challenge induces an inhibition of translational and transcriptional machinery

To understand the functional effects of the nutrient challenge, we performed a functional enrichment firstly on the set of downregulated genes defined above. We find that the set is enriched for translation-related genes (Figure 2C, SI Table 3). The most highly enriched terms in the Reactome annotation are related to translation, including ribosomal proteins, mitochondrial ribosomal proteins, translation initiation factors, and elongation factors. When examining the levels of all differentially expressed translation genes (annotated in Reactome), we see that on the whole, translation is downregulated as a result of nutrient intake, both at baseline and after the intervention (Figure 3A, SI Figure 3). Transcription, including mRNA splicing functionality is also enriched, together with transcription factors (Materials and Methods, p < 0.05, chi2 approximation to fisher exact test). Additionally, we find functionality related to the degradation and metabolism of RNA enriched in the set of downregulated genes.

### Nutrient challenge induces an immune activation and a stress response

The nutrient challenge, both at baseline and after the intervention, provokes a strong upregulation of genes enriched for innate immune functionality (Figure 2D, SI Table 3). The set is particularly enriched for the sub-annotation of Neutrophil degranulation. Many key immune genes, such as CD14, TLR4, NLRP3 and IL6R, are upregulated in response to the nutrient challenge (Figure 3B, SI Figure 3) e.g. Toll-like-Receptors, interleukin receptors, and Tumor Necrosis Factor Receptors are upregulated. Signal transduction is also enriched in the set of upregulated genes including STAT3. These observations, together with the observation that the blood cell type counts did not change (mixed effect model p > 0.05, SI Table 4), indicate an immune response and not a change in cell type composition. Specifically, an innate immune system response.

**Figure 3:**
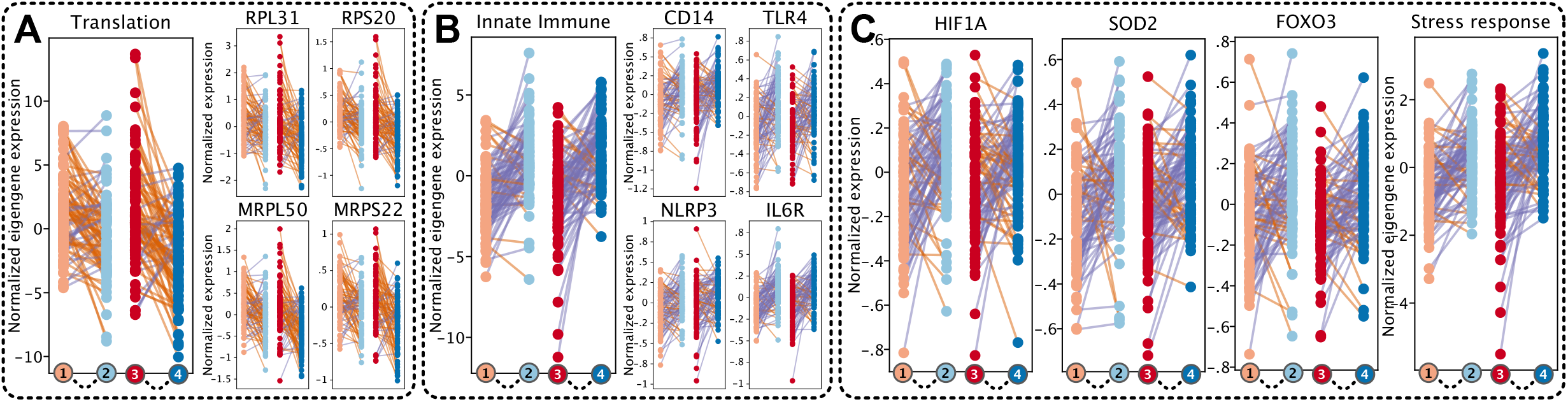
Per-individual gene responses to the nutrient challenge among specific gene groups. Each pair of points represents an individual’s response to the nutrient challenge, on the left at baseline, and on the right following the intervention. Each pair is connected with an orange line if the expression is increased, and with a blue line if the expression is decreased. In SI Figure 3, we provide these figures separately for males and females. A. For translation genes. From left-to-right: For the eigengene of all differentially expressed genes with the Reactome annotation of ‘Translation’, for RPL31, RPS20, MRPL50 and MRPS22.
B. For innate immune system genes. From left-to-right: For the eigengene of all differentially expressed genes with the Reactome annotation of ‘Innate immune system”, for CD14, TLR4, and IL6R.
C. For genes related to the regulation of cellular stress responses. From left-to-right: For HIF1A, SOD2, FOXO3, and all genes annotated in Reactome for ‘Cellular response to stress’.

In addition to the immune response, we also found that the cellular response to stress (Figure 2E, Reactome: R-HSA-2262752) was enriched. Among these, we see that FOXO3, HIF1A, TNF and SOD2 (Figure 3C, SI Figure 3) are significantly upregulated in response to the nutrient challenge, together with several heat shock protein transcripts.

### Intervention acts on the postprandial response

Previous results indicate an interaction between the nutrient challenge and the intervention, and we set out to investigate this effect. We visualize the interaction by plotting the trajectories of people in their nutrient response (Figure 4, SI Figure 4). We notice that the average trajectory is different between the baseline and post-intervention responses. The average fasted state is approximately the same at baseline and after the intervention (timepoints 1 and 3), both in males (Figure 4A) and females (Figure 4B), but the postprandial state (timepoints 2 and 4) is different, more so for males than females. This indicates that the intervention has modified the postprandial response.

**Figure 4:**
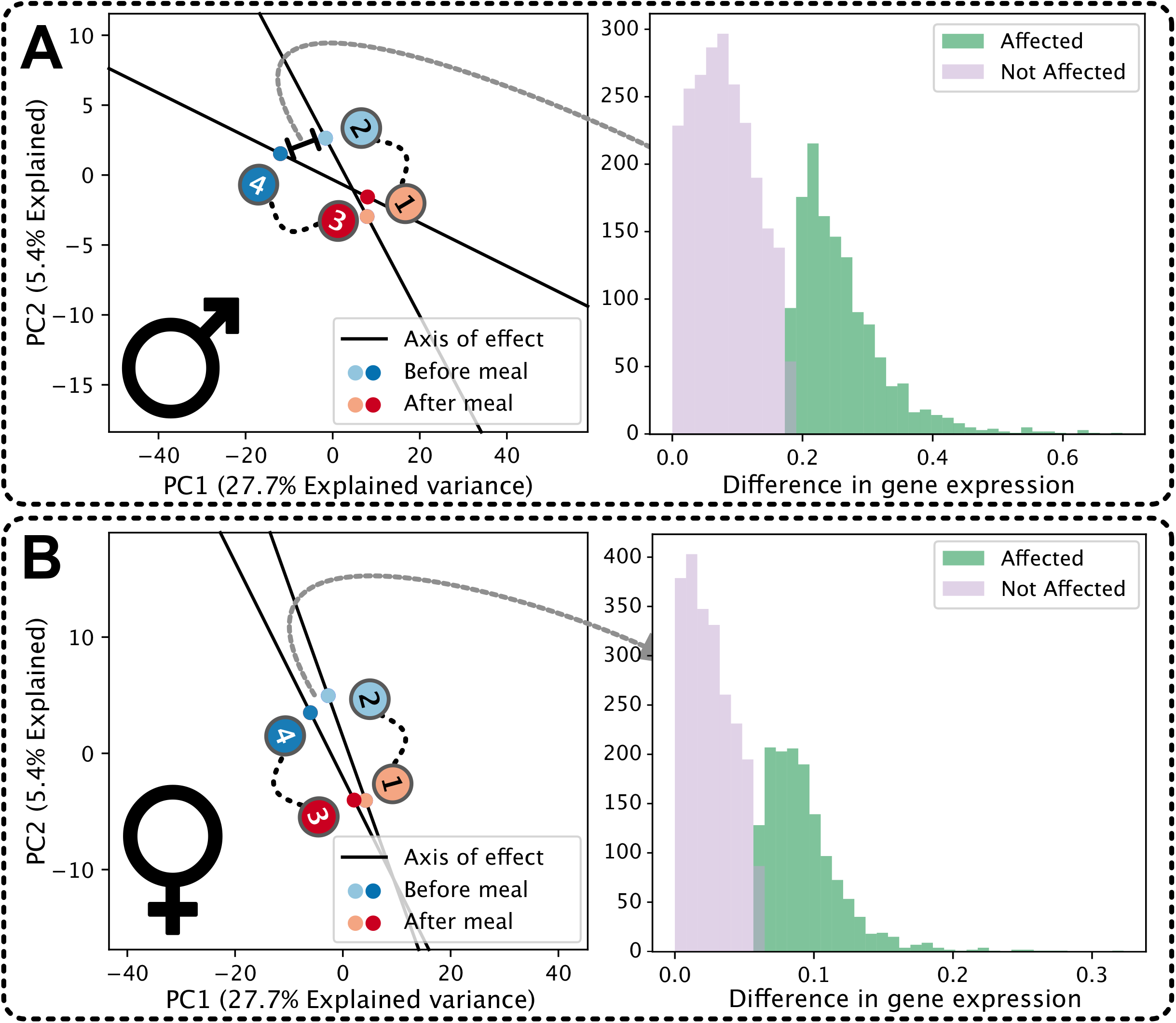
A. Trajectory analysis in PC space of males only.

1. Panel 1: Each point refers to a timepoint in the study (fasted in red, postprandial in blue), connected by a black line to indicate the nutrient challenge response trajectory. The baseline and post-intervention trajectories are superimposed to reveal that the intervention has rotated the postprandial response.
2. Panel 2: Projecting the postprandial trajectories back into gene space, we identify many genes whose postprandial response is lower expressed following the intervention.
B. As figure 4A, but for females only.

Upon formally testing this effect with an interaction model of differential expression, we found very few differentially expressed genes (44 in females, 129 in men, Methods, SI Table 2, SI Figure 1), despite the clear shift of trajectories in Figure 4, and the response in Figure 2A/B. No functional terms were enriched in these sets of genes. We hypothesized that the interaction effect between the nutrient challenge and the intervention was not much stronger than the nutrient challenge effect at baseline, that perhaps the heterogeneity between individuals was making the effect less clear.

Therefore, we determined the genes that contributed to the shift in trajectory in figures 4A and 4B (right panels, Materials and methods), finding 1413 genes in males, and 1304 in females, overlapping with 1175 genes. These genes were enriched for translational and transcriptional functions both in males and females (p < 0.05, fisher’s exact test, SI Table 3). This indicates that, although the intervention is acting differently on the magnitude of transcriptional response in males and females, it is acting on the same set of genes.

### Intervention tightens correlation network in response to the nutrient challenge across participants

To investigate the intervention induced effect further, we studied the gene-response correlation networks of the nutrient challenge response at baseline and after the intervention (Materials and methods). We investigated the correlation between gene responses for males and females separately. In both males and females at baseline, we find a high positive correlation between downregulated genes, and a high positive correlation (though not as high) between upregulated genes (for males: Figure 5A, first panel, for females: SI Figure 5). Between the up- and down-regulated genes we also see a high negative correlation, indicating that a number of people in our study have both the up− and down-regulation response, meaning that the magnitude of the response to the challenge is a characteristic of the individual. After the intervention (comparing timepoint 3 to 4), the correlations are more pronounced (for males: Figure 5A, second panel, for females: SI Figure 5). This indicates that the intervention reinforces the correlation between up- and down-regulated genes, that the intervention is modulating the postprandial response.

**Figure 5:**
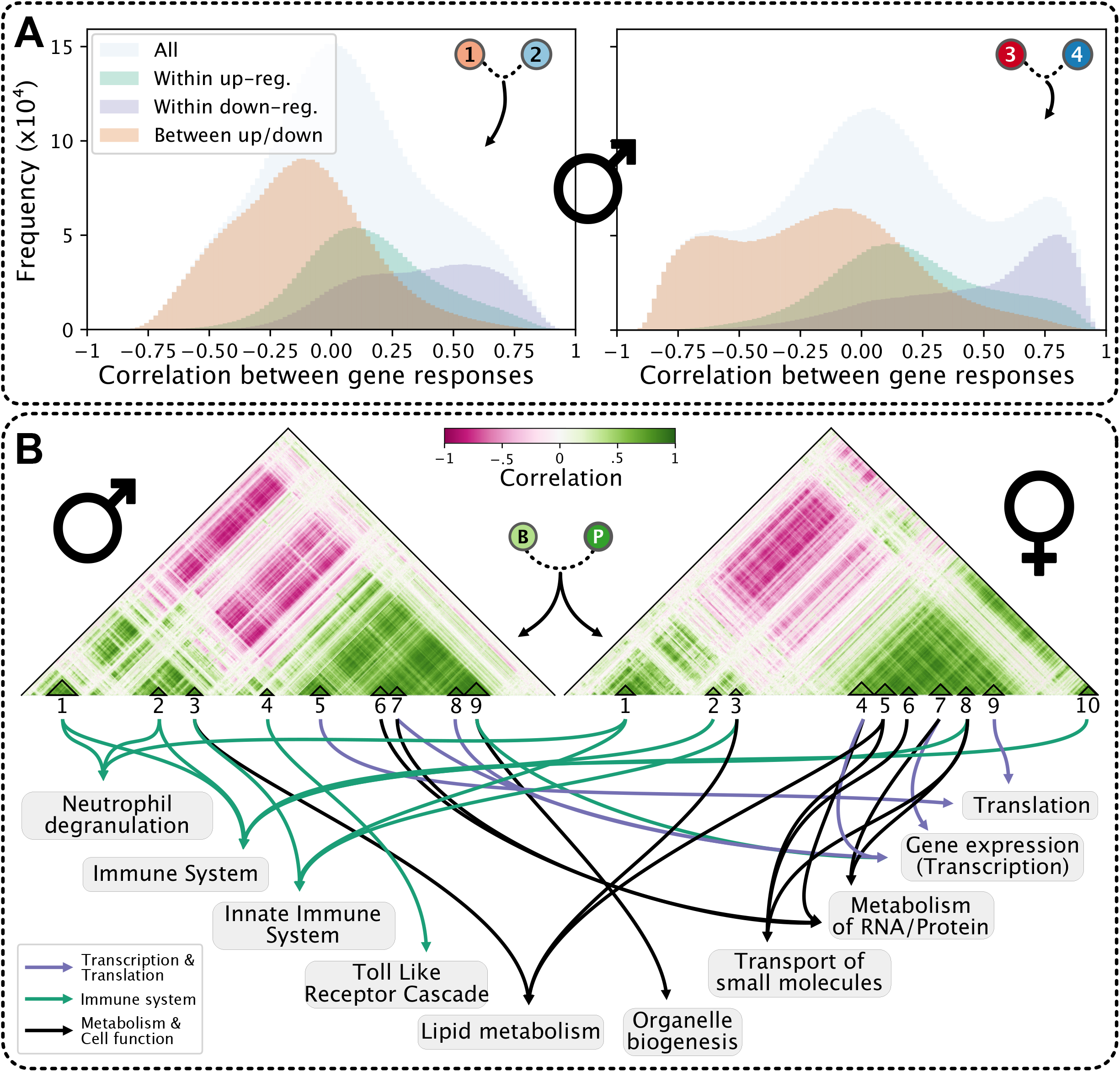
Modulation of the correlation network as a result of the intervention A. Modulation of the correlation network as a result of the intervention in males

1. Panel 1: The distribution of pairwise postprandial gene-response correlations at baseline. The green peak represents all correlations between up-regulated genes, the blue peak between all down-regulated genes, and the brown peak between up- and down-regulated genes.
2. Panel 2: The distribution of pairwise postprandial gene-response correlations following the intervention.
B. The correlation network between intervention-postprandial gene-responses. Green indicates a higher positive correlation, and red a higher negative correlation. The identified submodules are indicated, and arrows indicate the top functional enrichments for those clusters. The left network was generated using male gene responses only, and the right network using female responses only.

### Intervention induced changes in nutrient challenge based gene correlations differ between sexes

To better understand the change in correlation structure as a result of the intervention, we examined the correlation network of changes in the postprandial (i.e. how much stronger was the response at timepoint 3 to timepoint 4, versus timepoint 1 to timepoint 2) gene responses due to the intervention, separately for each sex (Materials and Methods). The correlation structures in the network are different for males and females (Figure 5B), indicating that the groups of genes that are modified by the intervention are different between the sexes. To understand the functional units of change in these networks, we detected modules in these networks. We identified 9 and 10 modules of highly correlated genes in males and females, respectively (Figure 5B, Materials and Methods). The clusters are composed of either up- or down-regulated genes. In males and females, clusters 1 to 3 are upregulated, and the remaining clusters (clusters 4 to 9 and 4 to 10 in males and females respectively) are downregulated. The down-regulated clusters are invariably related to translation and transcription and metabolism, and the up-regulated clusters are related to immune functionality (SI Table 3).

To examine the consequence of expression of these clusters with respect to health, we calculated the association between the detected modules and health indicators (separately in men and women). We did not find any significant association between the transcriptional levels of the gene modules in fasted blood and major physiological health parameters that changed as a result of the intervention such as BMI, systolic blood pressure, and fasted glucose levels. It is clear that the intervention is having an effect on the postprandial response (Figure 2), and since these do not associate with the physiological markers that changed with the intervention, it is likely that the changes are related to an intermediate effect that these general physiological markers do not capture.

### The metabolomic response to the challenge is largely unaffected by the intervention

Given that many studies have explored the response to challenge tests successfully by metabolomics analyses rather than transcriptome analyses, we recorded the response to the challenge by measuring 1H-NMR based blood metabolites at all four timepoints (timepoints 2 vs 1, and 4 vs 3, Figure 1). We calculated the sex-stratified effects of the nutrient challenge on the metabolites both before and after the intervention for individuals for whom we also have RNA-Sequencing data (Materials and Methods). As previously reported (Schutte et al., 2016), we find that the abundances of almost all metabolites are affected by the nutrient challenge (SI Figure 6). Additionally, we observe that the response to the nutrient challenge is consistent across the intervention; The directionality of the response is the same at baseline and after the intervention for the majority of the metabolites (SI Figure 2). Unlike the transcriptome, the intervention has little effect on the metabolomic response to the nutrient challenge. However, albumin and the percentage of Saturated Fatty Acids have different responses at baseline and after the intervention. Note that these effects are not the same for males and females, (Figure 6, SI Figure 6). Although these sex-differences are not significant (p>0.05, Materials and Methods), we observe that, both before and after the intervention, males have higher levels of glucose following the meal, and females have higher levels of leucine and isoleucine following the meal.

**Figure 6:**
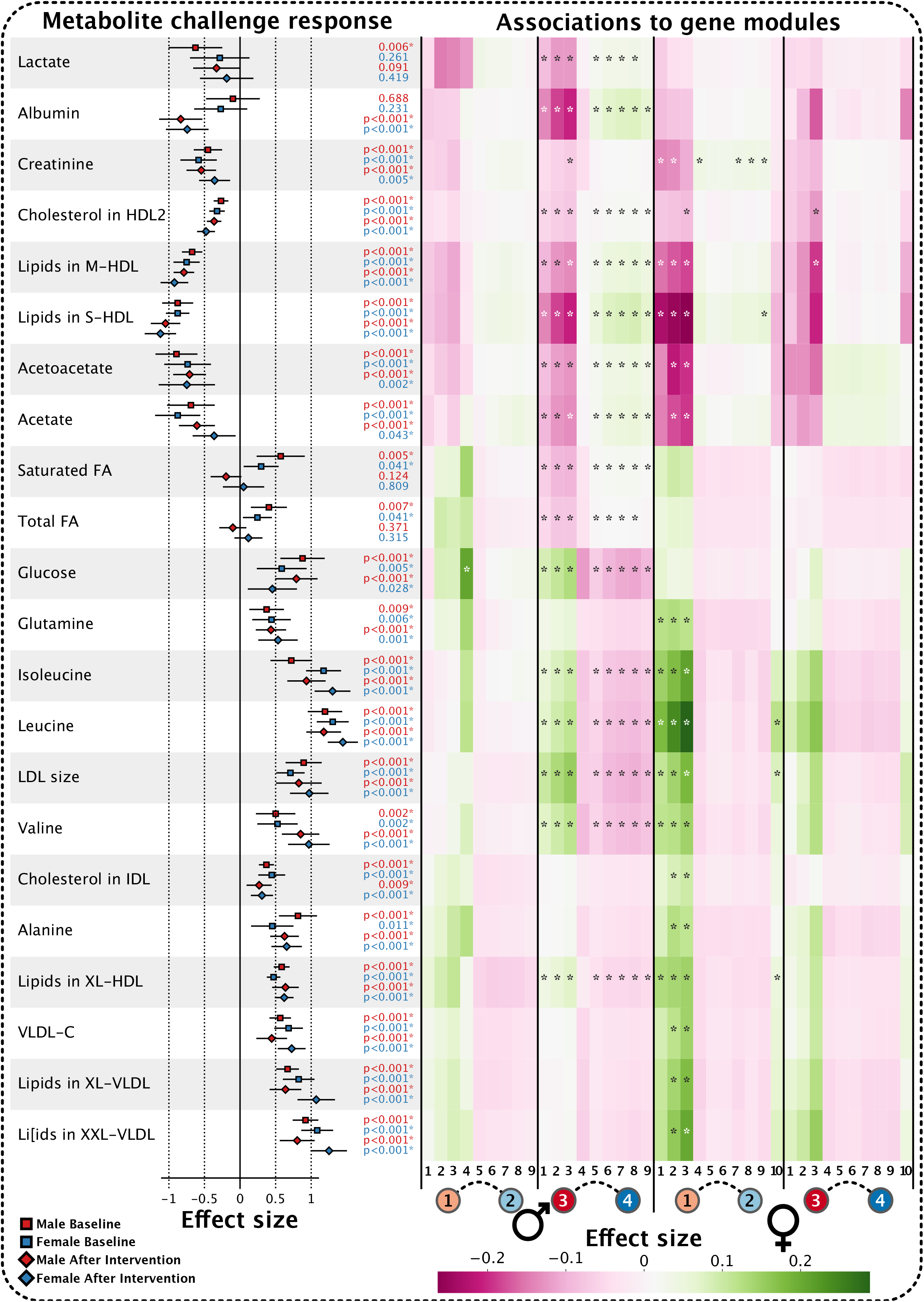
Metabolite changes as a result of the nutrient challenge. For each metabolite, on the left is shown a forest plot of the effect sizes of the nutrient challenge in males (in red, upper) females (in blue, lower) at baseline (square) and after the intervention (diamond). Error bars denote the 95% confidence interval. On the right, in the heatmap, the corresponding metabolite levels are associated with the gene module eigengenes under each condition, for males and females separately. From left to right: Men baseline, men after intervention, female baseline, female after intervention. An asterisk inside the heatmap corresponds to a significant association (p < 0.05). VLDL, very low density lipoprotein; LDL, low density lipoprotein; HDL, high density lipoprotein; FA, fatty acids; S, small; M, medium; L, large; XL; very large; XXL, extremely large.

### Metabolite levels are associated with gene modules during nutrient challenge

To understand the relationship between the metabolomic and transcriptomic responses to the nutrient challenge, and how these change as a result of the intervention, we investigated the association between the gene module eigengenes (representing a weighted average expression of genes in a specific module) and the metabolite levels, for males and females separately, at baseline, and after the intervention (Figure 6). We find that while the directionality of these associations are the same in males and females (for similar functionally-related modules), the strength of the associations differ, and are significant in different cases. The primary metabolite groups that associate with transcriptomic responses are glycolytic metabolites such as glucose, amino acids such as leucine and free fatty acids. At baseline, we find that the associations, both positive and negative between metabolites and the immune-related gene modules are stronger in females than in males. After the intervention, the opposite is true. This is consistent with our observations of the transcriptomic effects. Notable exceptions are glucose, which exhibits a stronger association with immune related gene modules at baseline than after the intervention in males, and none at all in females. For the translational modules (modules 5 and 6), we observe associations primarily in males after the intervention. Finally, the metabolomic results reveal the potential role of free fatty acids in the transcriptomic response, i.e. Albumin, Acetoacetate, Acetate, and Free Fatty acids are associated to the transcriptomic responses.

## Discussion

In the GOTO study, we subjected a population of healthy elderly Dutch individuals to a combined lifestyle intervention. Here we investigated to what extent a standardized nutrient challenge would reveal an individual molecular response to the intervention, that may go unnoticed in the traditionally sampled fasted blood. We observed that the intervention appeared to affect neither the fasted nor the postprandial (following an overnight fast) states of the blood transcriptome nor the postprandial state of the metabolome. The nutrient challenge itself elicited overall a strong consistent transcriptomic and metabolomic response. The transcriptomic response to the nutrient challenge at baseline and after the intervention was consistent for both sexes in terms of the responding genes and their directionality of change. This response was composed primarily of 1) an inhibition of ribosomal protein transcription, and 2) the activation of an innate immune response. The intervention affected this response, specifically on the ribosomal inhibition. Interestingly, whereas in males the response became stronger after the intervention, women had a stronger response at baseline. Further, the intervention increased the nutrient challenge based correlation between genes across participants, particularly in submodules related to the primary nutrient challenge response genes. Both the response to the challenge and modulation of the postprandial response by the intervention seemed a characteristic of the individual. After relating the gene modules of the transcriptomic response to metabolomic changes during the nutrient challenge, we found among sex differences, specific metabolites (glucose, specific amino acids, such as leucine, and free fatty acids) that correlate with the gene modules.

### Nutrient challenge stimulates a cellular stress response likely due to mitochondrial ROS

Following the nutrient challenge, glucose levels rise in the blood, both in males and females (Figure 6). Glucose stimulates the production of mitochondrial ROS in leukocytes (Mohanty et al., 2000), and we observe several downstream transcriptional responses that are indicative of this, including SOD2 (Hagenbuchner and Ausserlechner, 2013), HIF1A (Wellen and Thompson, 2010) and FOXO3 (Bakker et al., 2007; Kops et al., 2002). This constitutes a cellular stress response, and we hypothesize that the two major transcriptional effects we observe, the translational downregulation, and the innate immune activation, are components of this response. An impression of this is shown in Figure 7. As one of the most energy demanding tasks of a cell, translation is often the first to be modulated in a stress situation (Liu and Qian, 2014; Shenton et al., 2006; Spriggs et al., 2010). Similarly, the relationship between ROS and the innate immune system are well documented (Blaser et al., 2016; Chen et al., 2018; Park et al., 2015; Próchnicki and Latz, 2017; West et al., 2011), and immune responses to nutrient intake have previously been observed in blood (Baig et al., 2019; Leonardson et al., 2009). Our observations are consistent with an increased mitochondrial ROS production. Given that this response is occurring in 30 minutes following the meal, and that the participants are healthy, we expect that this is a normal, non-apoptotic, acute response to a change in homeostasis. We further stress that the overnight fast preceding the meal (‘refeeding’), may play a part in the biological responses we observed; The standardized challenge we have applied in effect is a liquid meal after an overnight fast. The composition of the standardized meal (SI Table 5) resembles a regular meal intake, however the 14 hours of fasting preceding the meal may be less regular.

**Figure 7:**
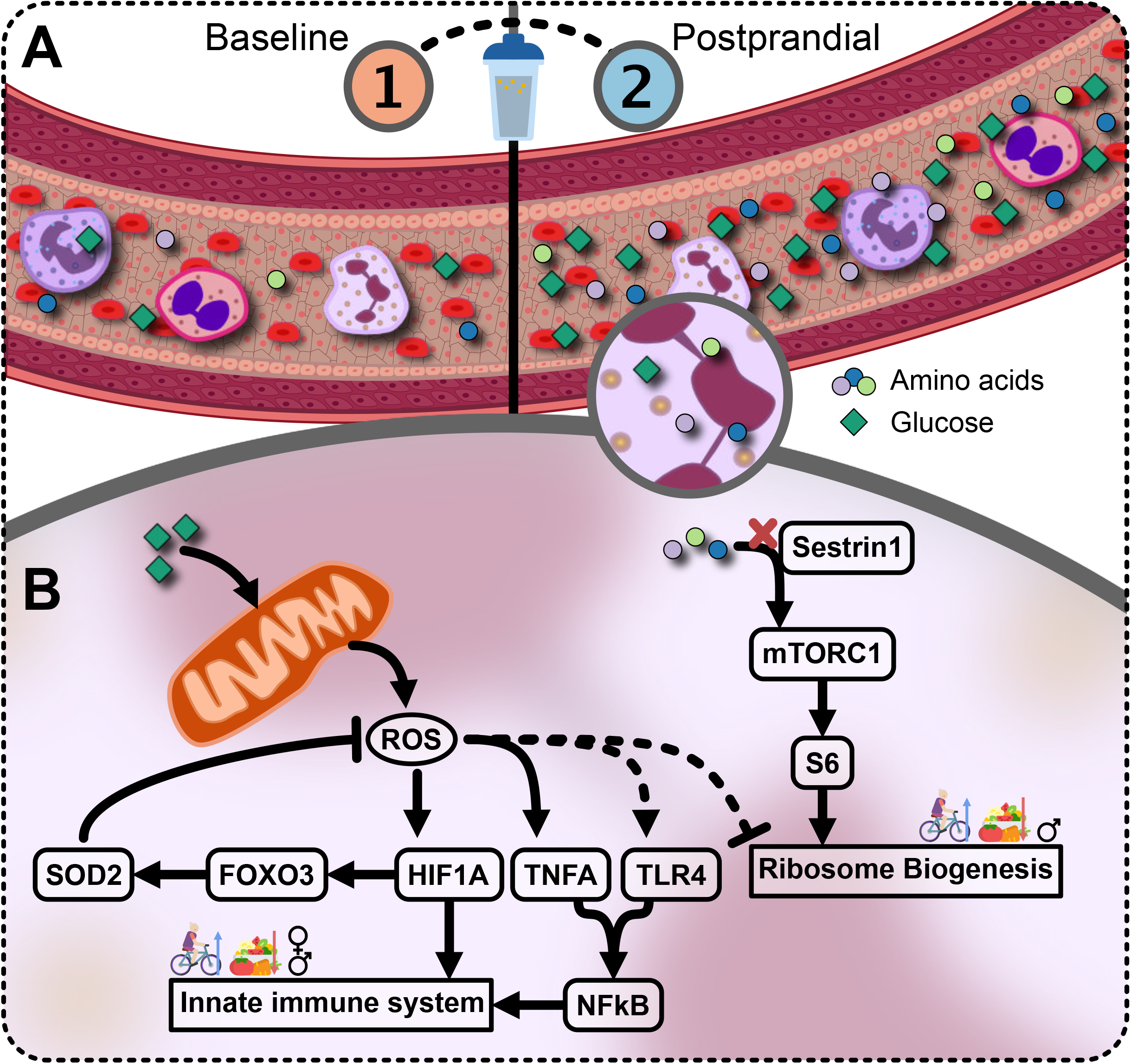
Impression of postprandial response

A. In the fasted state, the cells are in homeostasis. Nutrients are present, but slowly released into the bloodstream by the system. Following a meal, nutrients such as glucose and amino acids are now highly abundant, and are taken up by cells in the blood.
B. This abundance of glucose results in the activation of glycolysis and ROS production by mitochondria, which initiates a cascade resulting in the activation of innate immune system machinery. The amino acids which are taken up by the cells do not appear to stimulate mTORC1, and it may be possible that ROS, or other stress factors inhibit the activation of mTORC, and degradation of ribosomal protein transcript RNA, resulting in the downregulation of ribosome biosynthesis. These two effects are mediated by the intervention, and are sex specific.

### The lifestyle intervention sensitizes the system response to nutrient intake

The response to nutrient intake is consistent across the intervention, however, the intervention clearly affects the gene responses to nutrient intake. We observe not only that genes which affect the change in the postprandial response are primarily translational and immune genes, but also that the response correlation networks become tighter, and that specifically, the correlation within and between clusters relating to transcription, translation and immune function are being stimulated and strengthened. Together, this indicates that the translational and immune responses of the blood transcriptomic response to the nutrient challenge is being affected by the intervention.

To investigate the cause of the change in response to the nutrient challenge due to the intervention, we looked at the metabolomic response to the nutrient challenge. With the exception of albumin, there are no substantial changes in the metabolomic response at baseline vs after the intervention that would explain a different transcriptional response. Thus, the changes in the metabolomic environment of leukocytes due to the nutrient challenge has not substantially changed as a result of the intervention. Therefore, it must be the transcriptomic response to the metabolomic environment that changes. This is supported by the changes in the strengths of the association between metabolite and gene modules at baseline vs after the intervention, both in males and females. However, exactly how this is modulated is unclear. Although it is known that hypoalbuminemia is associated with increased levels of inflammation and TNFA (Gatta et al., 2012), it is not clear whether this relationship extends to the acute response we studied.

### The intervention modulates the postprandial response in a sex-dependent manner

When looking only at the sets of genes that we consider differentially expressed, it appears as though males and females have quite different transcriptional responses to the nutrient challenge. We have shown that this is not the case, and that the fundamental responses, a translational inhibition and an immune stimulation, are present in both sexes, merely at different magnitudes. Nevertheless, we did observe substantial differences between males and females in terms of the differential expression response to the nutrient challenge. Especially the translational element of the response, which was so pronounced following the intervention among bulk analysis is not found significant in females, although the directionality of the effect is consistent. Examining the gene responses to the intervention, we observe the same gene correlation structure in both sexes. Additionally, the network structure changes similarly in the two sexes as a result of the intervention. Thus, although the intervention affects the same nutrient challenge response in males and females, the direction of the effect is different in males and females. When looking at the serum metabolomics, we found that glucose behaves differently in males and females in response to the nutrient challenge, which may explain a difference in transcriptomic responses. On the other hand, for the most part, the metabolomic response to the challenge remains unchanged due to the intervention, but the association between metabolites and the gene modules do change considerably, and differently between the sexes. This indicates that, rather than changing the metabolomic response to the nutrient challenge, both sexes modulate their blood transcriptomic responses, but in different ways. It has been shown that men will consume more food if a meal is preceded by a fast, whereas a fast will induce women to consume less (Zandian et al., 2011). Whether this behavior is induced by social construction or hormonal signals in response to food is unclear, but it is possible that such biological effects play a role in our experiment also.

### Implications for health

We assume that the intervention has made people healthier. We have shown that the intervention has improved, among other things, the systolic blood pressure, body fat percentage, BMI and cholesterol levels of the subjects (van de Rest *et al.*, 2016, SI Table 1). To what extent do these systemic indicators of health reflect metabolic or cellular health? We investigated the association between the gene response submodules and physiological/health parameters, but did not find such associations for the gene modules of the translational response. The established physiological parameters are perhaps not able to capture these dynamics in the transcriptomic response to the nutrient challenge, or its alteration by the intervention. Hence, novel markers are needed to capture these cellular effects. This is complicated by the sex-specific results we observe. In females, the intervention results in a less pronounced, whereas in males it results in a more pronounced response to the nutrient challenge. It is possible that one sex is having a negative health response to the intervention, or alternatively that health may imply different cellular behavior in each sex?

We find that the transcriptome is more sensitive to these intervention effects than the metabolome, and may capture the phenotypic flexibility of the host’s stress response. Furthermore, the challenge we administered was not a diet stress test, such as high protein or lipid intake, but rather a standardized meal. Nevertheless, we observe a stress response, indicating that strong, stressful challenges may not be necessary to stimulate a measure of phenotypic flexibility. Our temporal resolution is limited to two samples within a 30 minute timeframe. With a higher resolution response over a longer period of time, we could better capture the dynamics of the intensity of the response, and duration until normalization. However, already with these two timepoints we have identified the stress response, and a have unveiled the interventions effect upon it. Further research on the dynamics of this response may lead to a better understanding of the suitability of this response in the search for a biomarker of phenotypic flexibility and health.

### Summary

In this work, we report on an acute, cellular response to fasting and refeeding that is sex- and individual-dependent, and which can be modulated by a combined physical exercise and nutritional intervention within 13 weeks. We note that the nutrient challenge was an essential component in understanding the cellular response to the intervention more so than the response of the metabolome. Without the challenge, we would not observe a transcriptomic change as a result of the intervention. The current practice is to investigate only fasted samples, but this may be limiting. Important homeodynamic effects from interventions may have gone unnoticed merely as a result of looking at an uninformative physiological state.

## Materials and Methods

### Data Availability

RNA-Seq counts and the nightingale metabolomics data will be made available upon request via the European Genome-Phenome Archive (EGA).

### Study design

In the Growing Old Together (GOTO) study, 164 (83 males, 81 females) healthy (mean BMI: 26.9 ± 2.5, no diagnosed inherited or metabolic diseases), elderly (mean age 62.9 ± 5.7) participants underwent a 13-week lifestyle intervention, as previously described (van de Rest et al., 2016; Schutte et al., 2016). The intervention consisted of a 25% change in energy balance, divided equally over an increased exercise regimen, and a decreased nutrient intake. Both at baseline (before) and after the intervention, we subjected participants to a nutrient challenge. This challenge is composed of an overnight fast and a subsequent refeeding with a standardized meal replacement shake. The nutridrink is a liquid oral nutritional supplement (Nutricia Advanced Medical Nutrition, Zoetermeer, The Netherlands; 1.5 kcal/mL (6.25 kJ/mL), 35% fat, 49% carbohydrates, 16% protein, SI Table 5). In a fasted and postprandial state (30 minutes after nutridrink administration), both at baseline and after the intervention, blood was sampled. Cell type counts were measured by Differential. Clinical chemistry parameters were measured in fasted serum collected by venipuncture. Of the 164 individuals in the GOTO study, we sequenced RNA of whole blood from 85 individuals (mean BMI: 26.8 ± 2.4, mean age: 63.2 ± 5.7, 44 males, 41 females) at all 4 time points. It is in these individuals in which we investigate our effects.

### Investigation of Intervention effects on health parameters

To test the effect of the intervention on measured health parameters (SI Table 1), we performed a mixed model test with a fixed effect of the intervention, and a random effect of the individual, both for a combined test including men and women, and a stratified test: *~ intervention + (1|person)*. P-values were adjusted for multiple testing with the Benjamini-Hochberg procedure to control for the False Discovery Rate.

### RNA isolation and sequencing

Libraries were prepared from whole blood RNA with Illumina TruSeq version 2 library preparation kits. With the Illumina HiSeq 2000 platform, paired-end sequencing reads (2 × 50-basepairs) were generated, with ten pooled samples per lane. Data processing was performed using the in-house BIOPET Gentrap pipeline, as previously described (Zhernakova et al., 2017). In short, low quality trimming was performed using sickle version 1.200 (‘se’ ‘−t’ ‘sanger’). Adapter clipping was performed using cutadapt version 1.1 (‘−m’ ‘25’). Reads were aligned to GRCh37, while masking common SNVs in the Dutch population (GoNL (Francioli et al., 2014) MAF>0.01), using STAR version 2.3.0e (‘-- outSAMstrandField’ ‘intronMotif’ ‘-- outSAMunmapped’ ‘Within’ ‘--outFilterMultimapNmax’ ‘5’ ‘--outFilterMismatchNmax’ ‘8’). Sam to bam conversion and sorting was performed using Picard version 2.4.1. Read quantification was performed using htseq-count version 0.6.1p1 (‘--format’ ‘bam’ ‘--order’ ‘pos’ ‘--stranded’ ‘no’) using Ensembl gene annotations version 71 for gene definitions. The sequencing resulted in an average of 37.1 million reads per sample, and 97.0 % (± 0.5 %) were mapped.

### Differential Gene Expression

There were 85 individuals for whom we have RNA-Seq data at all 4 timepoints. We identified confounding effects by testing the association between covariates and the principal components of the TMM-CPM normalized gene expression. We adjusted for technical effects (RNA isolation group, total μg yield, flowcell, mean insert size, and median 5’ bias), blood cell type percentages (eosinophils, monocytes, lymphocytes, basophil, red blood) and personal details (age, gender and individual, where individual was a random effect). In the interaction model, we added an interaction term between the intervention and fasted status. We removed genes that did not have at least 2 counts per sample on average. Differential expression was tested using limma with TMM and VOOM normalization (Law et al., 2014). We adjusted for multiple testing by correcting P-values with Bonferroni to control the Family-wise error rate. To select differentially expressed genes, a log2 fold change of 0.25 was applied. For the sex-stratified analysis, the same model was used, except without adjusting for sex. Due to the additional number of tests, we used the Benjamini-Hochberg procedure to correct for multiple testing across these conditions.

### Correlation network and module analysis

To adjust for the same effects as in the differential expression test, we fit a mixed model (with the same parameters as in the differential gene expression analysis) for each gene, and took the residual as the adjusted gene expression of the TMM-CPM expression values per person. For all genes that were differentially expressed between any of the conditions we tested, we calculated pearson correlation coefficients of the differences between the gene responses. Gene responses were calculated as the difference (fraction in log space) of the adjusted gene expression level before meal, and after the meal (after meal – before meal). These correlation networks were calculated both before and after the intervention. In this network, high correlations thus represent genes that change their food response due to the intervention in similar manners.

We determine these correlation networks of all genes that were differentially expressed in the nutrient challenge in either men or women. We calculated distances from the correlation matrix with D = 1-*p*, where *p* is the pearson correlation coefficient matrix. We performed hierarchical clustering on this distance matrix with complete linkage. Cutting the tree at a specific height was not appropriate, as the density of the nodes was different in different branches. Therefore, we calculated the modularity of clusters at each node in the tree using the modularity metric defined in (Ayroles et al., 2009), with an additional weighting factor of the height of the dendrogram at that node in the tree. We exclude clusters in which there are negative correlations between nodes (i.e. a height in the dendrogram above 1). We cut the tree based on the 95^th^ percentile modularity score, choosing a parent over its children only if both child nodes contained less than 100 genes. The dendrograms, and the clusters identified within the tree can be seen in SI Figure 7.

For each module, eigengenes were calculated, and multiplied by the sign of the correlation to the gene in the module which showed the highest absolute correlation with the eigengene; The eigengene set to be positively associated with the gene.

### Functional Enrichment

Functional enrichment was done with Fisher’s exact/chi^2^ approximation test on the Reactome (Fabregat et al., 2018) database, using the set of genes tested in the differential expression as a background. Tests were corrected using Benjamini Hochberg correction. For transcription factors, we used the list of transcription factors from Lambert et. al.’s review paper (Lambert et al., 2018). For the tests performed in the context of the PC trajectory analysis, and the context of the network modules, we used the background of genes used in that analysis. Figures 2D and 2E were generated by calculating a distance matrix between reactome terms with the following distance function, related to the inverse of the overlap between the genes annotated with two terms *s_i_* and 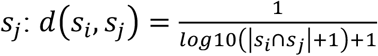. Terms were embedded in a dimensional space using multidimensional scaling. Terms with a distance less than 0.35 are connected by a line. Clustering was performed using complete hierarchical clustering on the distance matrix.

### Principal Component Trajectory Analysis

We adjusted for unwanted effects in the same way as for the correlation network analysis. Using again only the significant genes, we performed a PCA embedding of all samples. For each condition (before/after meal x before/after intervention), we calculated the centroid of all samples in the condition. We selected principal components which have a significant association (p<0.05) to the nutrient intake effect (linear model: principal component ~ effect). Using these components, we projected the centroids back into the original gene space. We calculated the difference between the back-projected centroids across the intervention condition, to find the difference gene expression between the fasted and activated samples. As we observed a bimodal distribution of differences (one near zero, and one with negative differences), we fitted a mixture model of two gaussians to the differences, and calculated the probability of each gene under the two distributions. We selected all genes which have a higher probability of not originating from the near zero distribution as the genes which have affected the food intake trajectory following the intervention.

### Nightingale metabolomics data

Using the serum of fasted and postprandial samples, both at baseline, and after the intervention, we quantified metabolomics using the nightingale 1-NMR metabolomics platform, as described in (Soininen et al., 2015). The effects of the intervention and nutrient challenge on the metabolome have previously been reported (van de Rest et al., 2016; Schutte et al., 2016). Of all individuals for whom we have metabolomics available at all four time points, 82 were overlapping with those for whom we have RNA-Seq data at all four time points. Of the 233 measures provided by the nightingale platform, several are derived measures, and we analysed the set of 63 non-derived metabolites (SI Table 6).

### Metabolomic analysis

Very little of the measured data (0.6%) were missing and were imputed using a recursive 3-nearest-neighbour system, in which missing values are taken as the average of the three nearest neighbors, determined by a euclidean distance to all other samples based on the non-missing metabolites for that sample. When all metabolites are imputed, distances can be recalculated and metabolites can be updated. This proceeds until the imputation has stabilized. Imputed values were all near zero (reflecting that missing values are near the detection limit). Metabolomic measures were log-transformed, (adding a pseudocount equal to the smallest non-zero metabolite measure), z-transformed and Rank Inverse Normal transformed. The effects of the nutrient challenge were investigated in a linear mixed model containing fixed effects for nutrient intake status, age, partner/offspring status, and blood cell type counts, and random effects for the individual, and household. Sex differences between the male and female responses the challenge were investigated at baseline and after the intervention with a linear mixed model with an interaction term between sex and nutrient intake status, with fixed effects for age, partner/offspring status, and blood cell type counts, and random effects for the individual, and household.

### Associating metabolites and gene modules

We investigated the association of a metabolite and the eigengene as they change across the nutrient challenge. Transformed metabolite levels and module eigengenes were investigated in a linear mixed model as *m ~ e + 1|person*, where *m* is a metabolite and *e* is a cluster eigengene. Associations were tested across the nutrient challenge within the baseline or post-intervention states.

## Data availability

Transcriptomic, metabolomic and associated metadata will be made available on the European Genome-Phenome Archive (EGA).

## ACKNOWLEDGEMENTS

The participants of our study, who adhered to the intervention, generously underwent testing and biopsy samples, and made this work possible.

