## Supplementary material for "A combined lifestyle intervention induces a sensitization of the blood transcriptomic response to a nutrient challenge": SI Figures and Tables

### Supplementary notes for “A combined lifestyle intervention induces a sensitization of the blood transcriptomic response to a nutrient challenge”

### Index

Index ... 2

SI Table 1: Effects of the intervention on parameters of body composition, health and functioning, and diagnostic measurements. ... 3

SI Table 2: Differentially expressed genes ... 6

SI Table 3: Functional Enrichments in different sets ... 6

SI Table 4: Blood cell type counts ... 7

SI Table 5: Comparison between the HuMet Standard Liquid Diet and the GOTO nutridrink ... 8

SI Table 6: Metabolites tested ... 9

SI Figure 1: Volcanoplots for combined and sex-stratified tests ... 12

SI Figure 2: Consistency of responses ... 14

SI Figure 3: Stratified Gene expression levels ... 16

SI Figure 4: Individual trajectories in PC Space ... 18

SI Figure 5: Change in correlation structure in males and females ... 20

SI Figure 6: Changes in metabolite levels by nutrient challenge ... 22

SI Figure 7: Modules detected in differentially expressed genes ... 24

Supplementary Citations ... 26

#### SI Table 1: Effects of the intervention on parameters of body composition, health and functioning, and diagnostic measurements.

We tested the effects of the intervention in our selection on all parameters that were significantly affected by the intervention among all participants of the study (van de Rest *et al.*, 2016).

| Body Composition | Group | n | Estimate | Std. Error | Pr(>\|t\|) | padj |
|---|---|---|---|---|---|---|
| BMI (kg/$m^2$) | Both | 85 | -0.491484 | 0.031611 | 1.91E-26 | 6.30E-25 |
| BMI (kg/$m^2$) | Men | 44 | -0.44216 | 0.045998 | 2.82E-12 | 2.33E-11 |
| BMI (kg/$m^2$) | Women | 41 | -0.544418 | 0.042122 | 7.28E-16 | 1.20E-14 |
| Weight (kg) | Both | 85 | -0.372314 | 0.023232 | 2.83E-27 | 1.87E-25 |
| Weight (kg) | Men | 44 | -0.365158 | 0.036661 | 9.73E-13 | 1.07E-11 |
| Weight (kg) | Women | 41 | -0.379994 | 0.02822 | 1.91E-16 | 4.19E-15 |
| Waist circumference (cm) | Both | 85 | -0.527537 | 0.062773 | 9.40E-13 | 1.07E-11 |
| Waist circumference (cm) | Men | 44 | -0.478792 | 0.087542 | 2.15E-06 | 9.46E-06 |
| Waist circumference (cm) | Women | 41 | -0.579848 | 0.090456 | 1.25E-07 | 7.52E-07 |
| Fat free mass ($kg^2$) | Both | 85 | -0.219552 | 0.032452 | 3.78E-09 | 2.77E-08 |
| Fat free mass ($kg^2$) | Men | 44 | -0.236645 | 0.053683 | 9.31E-05 | 2.95E-04 |
| Fat free mass ($kg^2$) | Women | 41 | -0.194882 | 0.034862 | 3.38E-06 | 1.40E-05 |
| Body fat (%) | Both | 85 | -0.226337 | 0.033573 | 4.19E-09 | 2.77E-08 |
| Body fat (%) | Men | 44 | -0.222536 | 0.043525 | 1.15E-05 | 4.47E-05 |
| Body fat (%) | Women | 41 | -0.230587 | 0.051856 | 9.40E-05 | 2.95E-04 |

| Health and Functioning | Group | n | Estimate | Std. Error | Pr(>\|t\|) | padj |
|---|---|---|---|---|---|---|
| Systolic bp, (mm Hg)[a] | Both | 61 | -0.263422 | 0.078973 | 1.46E-03 | 3.58E-03 |
| Systolic bp, (mm Hg)[a] | Men | 35 | -0.16209 | 0.107189 | 1.40E-01 | 2.25E-01 |
| Systolic bp, (mm Hg)[a] | Women | 26 | -0.39983 | 0.113075 | 1.61E-03 | 3.67E-03 |
| Diastolic bp, (mm Hg)[a] | Both | 61 | -0.353295 | 0.095142 | 4.50E-04 | 1.24E-03 |
| Diastolic bp, (mm Hg)[a] | Men | 35 | -0.305263 | 0.128178 | 2.30E-02 | 4.46E-02 |
| Diastolic bp, (mm Hg)[a] | Women | 26 | -0.417955 | 0.143525 | 7.45E-03 | 1.54E-02 |
| REE (kcal/day) | Both | 85 | -0.392101 | 0.06718 | 1.57E-07 | 8.63E-07 |
| REE (kcal/day) | Men | 44 | -0.354197 | 0.097443 | 8.75E-04 | 2.22E-03 |
| REE (kcal/day) | Women | 41 | -0.408054 | 0.091904 | 8.95E-05 | 2.95E-04 |
| SPPB | Both | 85 | 0.154229 | 0.111779 | 1.71E-01 | 2.57E-01 |
| SPPB | Men | 44 | 0.198628 | 0.161234 | 2.25E-01 | 3.15E-01 |
| SPPB | Women | 41 | 0.106581 | 0.155874 | 4.98E-01 | 5.77E-01 |

| Diagnostic Measures | Group | n | Estimate | Std. Error | Pr(>\|t\|) | padj |
|---|---|---|---|---|---|---|
| Log(Insulin) (mU/L) | Both | 85 | -0.043549 | 0.075492 | 5.66E-01 | 6.44E-01 |
| Log(Insulin) (mU/L) | Men | 44 | -0.078906 | 0.112407 | 4.86E-01 | 5.73E-01 |
| Log(Insulin) (mU/L) | Women | 41 | -0.005605 | 0.100822 | 9.56E-01 | 9.56E-01 |
| Cholesterol (mmol/L)[b] | Both | 72 | -0.341297 | 0.096833 | 7.47E-04 | 1.97E-03 |
| Cholesterol (mmol/L)[b] | Men | 37 | -0.29807 | 0.140525 | 4.09E-02 | 7.29E-02 |

| | | | | | | |
|---|---|---|---|---|---|---|
| Cholesterol (mmol/L)[b] | Women | 35 | -0.386994 | 0.134375 | 6.84E-03 | 1.46E-02 |
| Cholesterol_HDL (mmol/L)[b] | Both | 72 | -0.06479 | 0.069006 | 3.51E-01 | 4.52E-01 |
| Cholesterol_HDL (mmol/L)[b] | Men | 37 | 0.156036 | 0.082284 | 6.60E-02 | 1.15E-01 |
| Cholesterol_HDL (mmol/L)[b] | Women | 35 | -0.298234 | 0.098785 | 4.78E-03 | 1.05E-02 |
| LDL_Cholesterol (mmol/L)[b] | Both | 72 | -0.342264 | 0.081633 | 7.83E-05 | 2.87E-04 |
| LDL_Cholesterol (mmol/L)[b] | Men | 37 | -0.414995 | 0.12158 | 1.60E-03 | 3.67E-03 |
| LDL_Cholesterol (mmol/L)[b] | Women | 35 | -0.265376 | 0.108364 | 1.96E-02 | 3.93E-02 |
| Log(triglycerides) (mmol/L)b | Both | 72 | 0.101197 | 0.046541 | 3.30E-02 | 6.23E-02 |
| Log(triglycerides) (mmol/L)b | Men | 37 | 0.265653 | 0.068206 | 4.09E-04 | 1.20E-03 |
| Log(triglycerides) (mmol/L)b | Women | 35 | -0.072656 | 0.048663 | 1.45E-01 | 2.27E-01 |
| TSH (mU/L)b | Both | 72 | -0.092626 | 0.069409 | 1.86E-01 | 2.67E-01 |
| TSH (mU/L)b | Men | 37 | -0.12726 | 0.093594 | 1.82E-01 | 2.67E-01 |
| TSH (mU/L)b | Women | 35 | -0.056013 | 0.103989 | 5.94E-01 | 6.53E-01 |
| DHEAS (nmol/L)c | Both | 85 | -0.031264 | 0.034452 | 3.67E-01 | 4.57E-01 |
| DHEAS (nmol/L)c | Men | 44 | 0.015244 | 0.046221 | 7.43E-01 | 7.91E-01 |
| DHEAS (nmol/L)c | Women | 41 | -0.081176 | 0.05082 | 1.18E-01 | 2.00E-01 |
| Log(Leptin) (µg/L) | Both | 85 | -0.373918 | 0.045596 | 2.40E-12 | 2.27E-11 |
| Log(Leptin) (µg/L) | Men | 44 | -0.413556 | 0.069387 | 4.18E-07 | 2.12E-06 |
| Log(Leptin) (µg/L) | Women | 41 | -0.331379 | 0.05839 | 1.35E-06 | 6.37E-06 |
| Log(Adiponectin) (mg/L) | Both | 61 | 0.11647 | 0.05465 | 3.72E-02 | 6.82E-02 |
| Log(Adiponectin) (mg/L) | Men | 35 | 0.263722 | 0.067419 | 4.17E-04 | 1.20E-03 |
| Log(Adiponectin) (mg/L) | Women | 26 | -0.081754 | 0.075869 | 2.92E-01 | 3.93E-01 |
| IGF1 (nmol/L) | Both | 85 | 0.024569 | 0.05319 | 6.45E-01 | 6.98E-01 |
| IGF1 (nmol/L) | Men | 44 | 0.057414 | 0.072316 | 4.32E-01 | 5.28E-01 |
| IGF1 (nmol/L) | Women | 41 | -0.01068 | 0.078894 | 8.93E-01 | 9.07E-01 |
| IGFBP-3 (mg/L) | Both | 85 | -0.079955 | 0.08051 | 3.24E-01 | 4.27E-01 |
| IGFBP-3 (mg/L) | Men | 44 | -0.176558 | 0.119957 | 1.48E-01 | 2.28E-01 |
| IGFBP-3 (mg/L) | Women | 41 | 0.023716 | 0.105355 | 8.23E-01 | 8.49E-01 |
| IGF-1:IGFBP-3 | Both | 85 | 0.099248 | 0.084379 | 2.43E-01 | 3.34E-01 |
| IGF-1:IGFBP-4 | Men | 44 | 0.21594 | 0.13713 | 1.23E-01 | 2.02E-01 |
| IGF-1:IGFBP-5 | Women | 41 | -0.025982 | 0.092444 | 7.80E-01 | 8.17E-01 |
| CRP (high-sensitivity), (mg/L)a | Both | 85 | -0.103727 | 0.1118 | 3.56E-01 | 4.52E-01 |
| CRP (high-sensitivity), (mg/L)a | Men | 44 | -0.085597 | 0.156706 | 5.88E-01 | 6.53E-01 |
| CRP (high-sensitivity), (mg/L)a | Women | 41 | -0.123183 | 0.161458 | 4.50E-01 | 5.40E-01 |
| fT3 (pmol/L) | All | 85 | -0.153412 | 0.034946 | 3.28E-05 | 8.20E-05 |
| fT3 (pmol/L) | Men | 44 | -0.155909 | 0.045026 | 1.22E-03 | 2.12E-03 |
| fT3 (pmol/L) | Women | 41 | -0.150732 | 0.054567 | 8.63E-03 | 1.14E-02 |

[a]Individuals on antihypertensive medication were excluded from this analysis
[b]Individuals on lipid-lowering medication were excluded from this analysis

#### SI Table 2: Differentially expressed genes

See SI_Table_2.xlsx for a list (and summary) of all differentially expressed genes

#### SI Table 3: Functional Enrichments in different sets

Supplementary file SI_Table_3.xlsx provides the significant terms identified in:

- The set of down-regulated genes in response to the nutrient challenge, before or after the intervention, in males or females
- The set of up-regulated genes in response to the nutrient challenge, before or after the intervention, in males or females
- The set of up or down regulated genes in response to the intervention in a fasted state
- The set of up or down regulated genes in response to the intervention in a postprandial state.
- The set of genes found to affect the trajectory of the nutrient challenge response in males
- The set of genes found to affect the trajectory of the nutrient challenge response in females
- The sets of genes in each module identified in the male correlation network
- The sets of genes in each module identified in the female correlation network

#### SI Table 4: Blood cell type counts

No blood cell type was differently represented between at baseline versus after the intervention ($p > 0.05$), also not in a sex-stratified analysis. Blood cell type count percentages were assessed with a linear mixed model with *percentage ~ intervention + sex + (1|person)* for the bulk analysis, and with ~ intervention + (1|person_id) for the sex-stratified analysis.

| Estimate | Std..Error | t.value | p-value | Celltype | Condition | q-value |
|---|---|---|---|---|---|---|
| 0.001534 | 0.001324 | 1.158072 | 2.49E-01 | hematocrit | All | 5.27E-01 |
| 0.008282 | 0.104856 | 0.078987 | 9.37E-01 | eosinophiles | All | 9.54E-01 |
| 0.031288 | 0.022545 | 1.387845 | 1.67E-01 | basophiles | All | 4.27E-01 |
| -0.028896 | 0.497674 | -0.058062 | 9.54E-01 | neutrophiles | All | 9.54E-01 |
| -0.168896 | 0.427137 | -0.395414 | 6.93E-01 | lymphocyte | All | 8.66E-01 |
| 0.163681 | 0.167841 | 0.975214 | 3.31E-01 | monocytes | All | 5.84E-01 |
| -0.000854 | 0.001895 | -0.450587 | 6.53E-01 | hematocrit | Male | 8.52E-01 |
| 0.097683 | 0.150023 | 0.651121 | 5.17E-01 | eosinophiles | Male | 8.16E-01 |
| 0.056951 | 0.037235 | 1.529527 | 1.30E-01 | basophiles | Male | 3.90E-01 |
| 0.345122 | 0.697494 | 0.494803 | 6.22E-01 | neutrophiles | Male | 8.48E-01 |
| -0.835732 | 0.604836 | -1.38175 | 1.71E-01 | lymphocyte | Male | 4.27E-01 |
| 0.342683 | 0.215171 | 1.592607 | 1.15E-01 | monocytes | Male | 3.84E-01 |
| 0.003951 | 0.001823 | 2.167065 | 3.32E-02 | hematocrit | Female | 1.73E-01 |
| -0.082222 | 0.146733 | -0.560353 | 5.77E-01 | eosinophiles | Female | 8.24E-01 |
| 0.005309 | 0.025166 | 0.210949 | 8.33E-01 | basophiles | Female | 9.54E-01 |
| -0.407531 | 0.712088 | -0.572304 | 5.69E-01 | neutrophiles | Female | 8.24E-01 |
| 0.506173 | 0.597637 | 0.846957 | 4.00E-01 | lymphocyte | Female | 6.66E-01 |
| -0.017531 | 0.257921 | -0.06797 | 9.46E-01 | monocytes | Female | 9.54E-01 |

#### SI Table 5: Comparison between the HuMet Standard Liquid Diet and the GOTO nutridrink

Caseine AA concentrations were taken from (Bjørndal *et al.*, 2015).

| **Component** | **GOTO Nutricia energy drink, vanilla** | **Humet Fresubin Energy Drink, chocolate** |
|---|---|---|
| Carbohydrates | 18.4 | 18.8 |
| glucose | 0.1 | 0.2 |
| fructose | 0 | 0 |
| maltose | 0.6 | 0.33 |
| sucrose | 6 | 5.5 |
| lactose | 0.025 | 0.27 |
| polysacc | 11.3 | 12.45 |
| Protein* | 5.9 | 6 |
| lysine | 0.4366 | 0.45 |
| threonine | 0.2242 | 0.25 |
| methionine | 0.1534 | 0.15 |
| phenylalanine | 0.295 | 0.28 |
| tryptophan | 0.0708 | 0.08 |
| valine | 0.3363 | 0.39 |
| leucine | 0.5192 | 0.54 |
| isoleucine | 0.2832 | 0.3 |
| tyrosone | 0.3127 | 0.3 |
| cysteine | 0.0236 | 0.004 |
| histidine | 0.1534 | 0.15 |
| arginine | 0.2124 | 0.21 |
| glutamine | | 0.47 |
| glycine | 0.1062 | 0.11 |
| alanine | 0.1534 | 0.18 |
| proline | 0.6903 | 0.54 |
| serine | 0.3186 | 0.32 |
| glutamic acid | 1.2272 | 0.71 |
| aspartic acid & asparagine | 0.3835 | 0.57 |

#### SI Table 6: Metabolites tested

| Abbreviation | Name | Group | Subgroup | Unit |
|---|---|---|---|---|
| Ala | Alanine | Amino acids | Amino acids | mmol/l |
| Gln | Glutamine | Amino acids | Amino acids | mmol/l |
| His | Histidine | Amino acids | Amino acids | mmol/l |
| Phe | Phenylalanine | Amino acids | Aromatic amino acids | mmol/l |
| Tyr | Tyrosine | Amino acids | Aromatic amino acids | mmol/l |
| Ile | Isoleucine | Amino acids | Branched-chain amino acids | mmol/l |
| Leu | Leucine | Amino acids | Branched-chain amino acids | mmol/l |
| Val | Valine | Amino acids | Branched-chain amino acids | mmol/l |
| ApoA1 | Apolipoprotein A-I | Apolipoproteins | Apolipoproteins | g/l |
| ApoB | Apolipoprotein B | Apolipoproteins | Apolipoproteins | g/l |
| Total-C | Total cholesterol | Cholesterol | Cholesterol | mmol/l |
| LDL-C | Total cholesterol in 3 LDL subclasses | Cholesterol | Cholesterol | mmol/l |
| HDL-C | Total cholesterol in HDL | Cholesterol | Cholesterol | mmol/l |
| HDL2-C | Total cholesterol in HDL2 | Cholesterol | Cholesterol | mmol/l |
| HDL3-C | Total cholesterol in HDL3 | Cholesterol | Cholesterol | mmol/l |
| VLDL-C | Total cholesterol in VLDL | Cholesterol | Cholesterol | mmol/l |
| MUFA % | Ratio of monounsaturated fatty acids to total fatty acids | Fatty acids | Fatty acid ratios | % |
| Omega-3 % | Ratio of omega-3 fatty acids to total fatty acids | Fatty acids | Fatty acid ratios | % |
| Omega-6 % | Ratio of omega-6 fatty acids to total fatty acids | Fatty acids | Fatty acid ratios | % |
| PUFA % | Ratio of polyunsaturated fatty acids to total ... | Fatty acids | Fatty acid ratios | % |
| SFA % | Ratio of saturated fatty acids to total fatty ... | Fatty acids | Fatty acid ratios | % |
| Unsaturation | Degree of unsaturation | Fatty acids | Fatty acids | degree |
| DHA | Docosahexaenoic acid; 22:6 | Fatty acids | Fatty acids | mmol/l |
| LA | Linoleic acid; 18:2 | Fatty acids | Fatty acids | mmol/l |

| | | | | |
|---|---|---|---|---|
| MUFA | Monounsaturated fatty acids; 16:1, 18:1 | Fatty acids | Fatty acids | mmol/l |
| Omega-3 | Omega-3 fatty acids | Fatty acids | Fatty acids | mmol/l |
| Omega-6 | Omega-6 fatty acids | Fatty acids | Fatty acids | mmol/l |
| PUFA | Polyunsaturated fatty acids | Fatty acids | Fatty acids | mmol/l |
| SFA | Saturated fatty acids | Fatty acids | Fatty acids | mmol/l |
| Total-FA | Total fatty acids | Fatty acids | Fatty acids | mmol/l |
| Albumin | Albumin | Fluid balance | Fluid balance | signal area |
| Creatinine | Creatinine | Fluid balance | Fluid balance | mmol/l |
| Phosphatidylc | Phosphatidylcholine | Glycerides and phospholipids | Glycerides and phospholipids | mmol/l |
| Sphingomyelins | Sphingomyelins | Glycerides and phospholipids | Glycerides and phospholipids | mmol/l |
| Cholines | Total cholines | Glycerides and phospholipids | Glycerides and phospholipids | mmol/l |
| Total-TG | Total triglycerides | Glycerides and phospholipids | Glycerides and phospholipids | mmol/l |
| Phosphoglyc | Total phosphoglycerides | Glycerides and phospholipids | Glycerides and phospholipids | mmol/l |
| Citrate | Citrate | Glycolysis related metabolites | Glycolysis related metabolites | mmol/l |
| Glucose | Glucose | Glycolysis related metabolites | Glycolysis related metabolites | mmol/l |
| Lactate | Lactate | Glycolysis related metabolites | Glycolysis related metabolites | mmol/l |
| Pyruvate | Pyruvate | Glycolysis related metabolites | Glycolysis related metabolites | mmol/l |
| GlycA | Glycoprotein acetyls (GlycA) | Inflammation | Inflammation | mmol/l |
| bOHbutyrate | 3-hydroxybutyrate | Ketone bodies | Ketone bodies | mmol/l |
| Acetate | Acetate | Ketone bodies | Ketone bodies | mmol/l |
| Acetoacetate | Acetoacetate | Ketone bodies | Ketone bodies | mmol/l |
| HDL size | Mean diameter for HDL particles | Lipoprotein particle sizes | Lipoprotein particle sizes | nm |
| LDL size | Mean diameter for LDL particles | Lipoprotein particle sizes | Lipoprotein particle sizes | nm |
| VLDL size | Mean diameter for VLDL particles | Lipoprotein particle sizes | Lipoprotein particle sizes | nm |
| XXL-VLDL-L | Total lipids in chylomicrons and extremely lar... | Lipoprotein subclasses | Chylomicrons and extremely large VLDL | mmol/l |

| | | | | |
|---|---|---|---|---|
| IDL-C | Total cholesterol in IDL | Lipoprotein subclasses | IDL | mmol/l |
| IDL-L | Total lipids in IDL | Lipoprotein subclasses | IDL | mmol/l |
| L-HDL-L | Total lipids in large HDL | Lipoprotein subclasses | Large HDL | mmol/l |
| L-LDL-L | Total lipids in large LDL | Lipoprotein subclasses | Large LDL | mmol/l |
| L-VLDL-L | Total lipids in large VLDL | Lipoprotein subclasses | Large VLDL | mmol/l |
| M-HDL-L | Total lipids in medium HDL | Lipoprotein subclasses | Medium HDL | mmol/l |
| M-LDL-L | Total lipids in medium LDL | Lipoprotein subclasses | Medium LDL | mmol/l |
| M-VLDL-L | Total lipids in medium VLDL | Lipoprotein subclasses | Medium VLDL | mmol/l |
| S-HDL-L | Total lipids in small HDL | Lipoprotein subclasses | Small HDL | mmol/l |
| S-LDL-L | Total lipids in small LDL | Lipoprotein subclasses | Small LDL | mmol/l |
| S-VLDL-L | Total lipids in small VLDL | Lipoprotein subclasses | Small VLDL | mmol/l |
| XL-HDL-L | Total lipids in very large HDL | Lipoprotein subclasses | Very large HDL | mmol/l |
| XL-VLDL-L | Total lipids in very large VLDL | Lipoprotein subclasses | Very large VLDL | mmol/l |
| XS-VLDL-L | Total lipids in very small VLDL | Lipoprotein subclasses | Very small VLDL | mmol/l |

#### SI Figure 1: Volcanoplots for combined and sex-stratified tests

Volcano plots for each differential expression test. Red lines indicate the FDR adjusted p-value threshold. Yellow lines indicate effect size thresholds. Green points represent significant genes, and purple points are not significant. In each panel, we provide the unstratified, as well as the sex stratified results. The number of genes differentially expressed are indicated in bold text.

A) Fasted intervention effect
B) Postprandial intervention effect. Note that interpreting the intervention effect in only the postprandial samples is problematic, as it ignores the context of the fasted starting point in terms of investigating the response. A proper investigation of this effect can be seen in the interaction test (given below).
C) Baseline challenge effect
D) Post-intervention challenge effect
E) Intervention*challenge Interaction effect

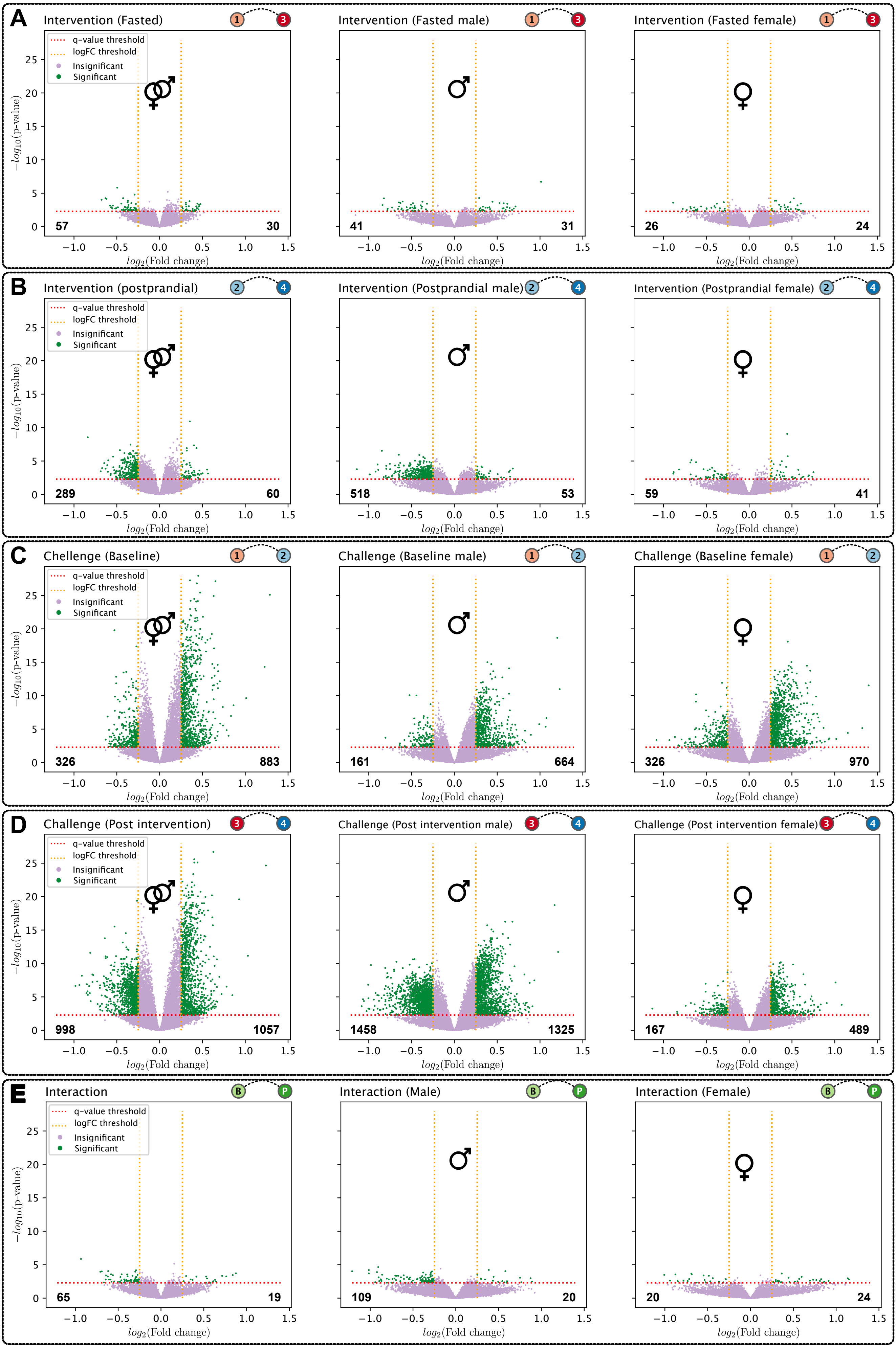

A
Intervention (Fasted)
Intervention (Fasted male)
Intervention (Fasted female)
57
30
41
31
26
24
B
Intervention (postprandial)
Intervention (Postprandial male)
Intervention (Postprandial female)
289
60
518
53
59
41
C
Chellenge (Baseline)
Challenge (Baseline male)
Challenge (Baseline female)
326
883
161
664
326
970
D
Challenge (Post intervention)
Challenge (Post intervention male)
Challenge (Post intervention female)
998
1057
1458
1325
167
489
E
Interaction
Interaction (Male)
Interaction (Female)
65
19
109
20
20
24
q-value threshold
logFC threshold
Insignificant
Significant
$-log_{10}$(p-value)
$log_2$(Fold change)

#### SI Figure 2: Consistency of responses

A) Consistency of responses between males and females in the transcriptomic nutrient challenge response. Provided as a paired-volcano plot. Green points are female gene differential expression test results, linked by a line to the male response in purple. Above, the baseline response, below, the post-intervention response.
B) Consistency of responses between males and females in the transcriptomic intervention response. Provided as a paired-volcano plot. Green points are female gene differential expression test results, linked by a line to the male response in purple. Above, the fasted response, below, the postprandial response.
C) Consistency of the metabolomic nutrient response. Provided as a paired-volcano plot. Left panel is for males, and right panel is for females. Blue points are baseline metabolite responses, red points, connected to blue points with a line are post intervention metabolomic responses.

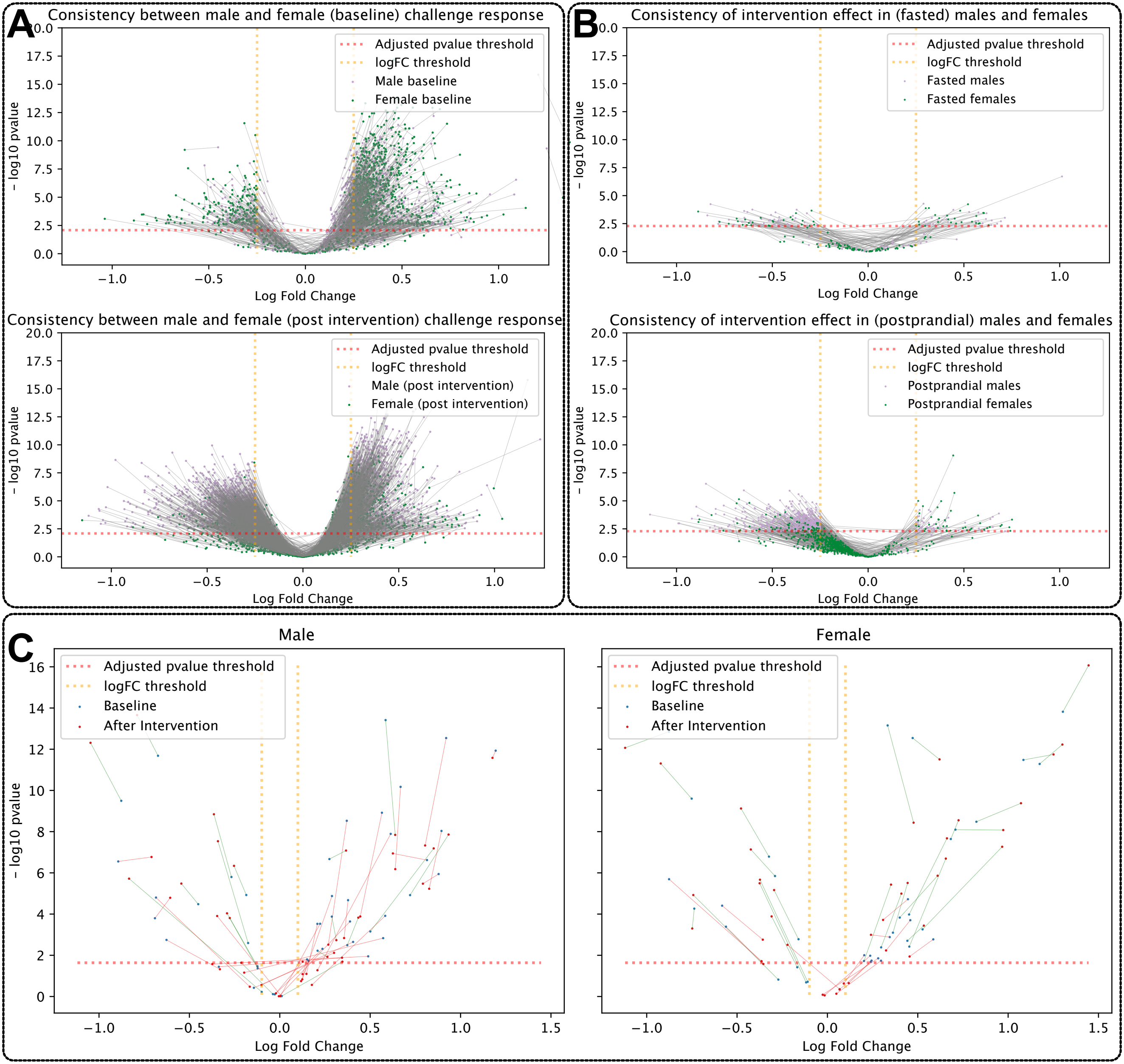
A
Consistency between male and female (baseline) challenge response
Adjusted pvalue threshold
logFC threshold
Male baseline
Female baseline
− log10 pvalue
Log Fold Change
Consistency between male and female (post intervention) challenge response
Male (post intervention)
Female (post intervention)
B
Consistency of intervention effect in (fasted) males and females
Fasted males
Fasted females
Consistency of intervention effect in (postprandial) males and females
Postprandial males
Postprandial females
C
Male
Female
Baseline
After Intervention

#### SI Figure 3: Stratified Gene expression levels

Expression levels shown in Figure 3 are for both males and females together. Here we provide the same plots as in Figure 3, but stratified for males and females, to show that the effects are consistent across the sexes, and differ in intensity. Each pair of points represents an individual's response to the nutrient challenge, on the left at baseline, and on the right following the intervention. Each pair is connected with an orange line if the expression is increased, and with a blue line if the expression is decreased. In SI Figure 3, we provide these figures separately for males and females.

- **A)** The translational genes
- **B)** The innate immune genes, and
- **C)** The stress genes.

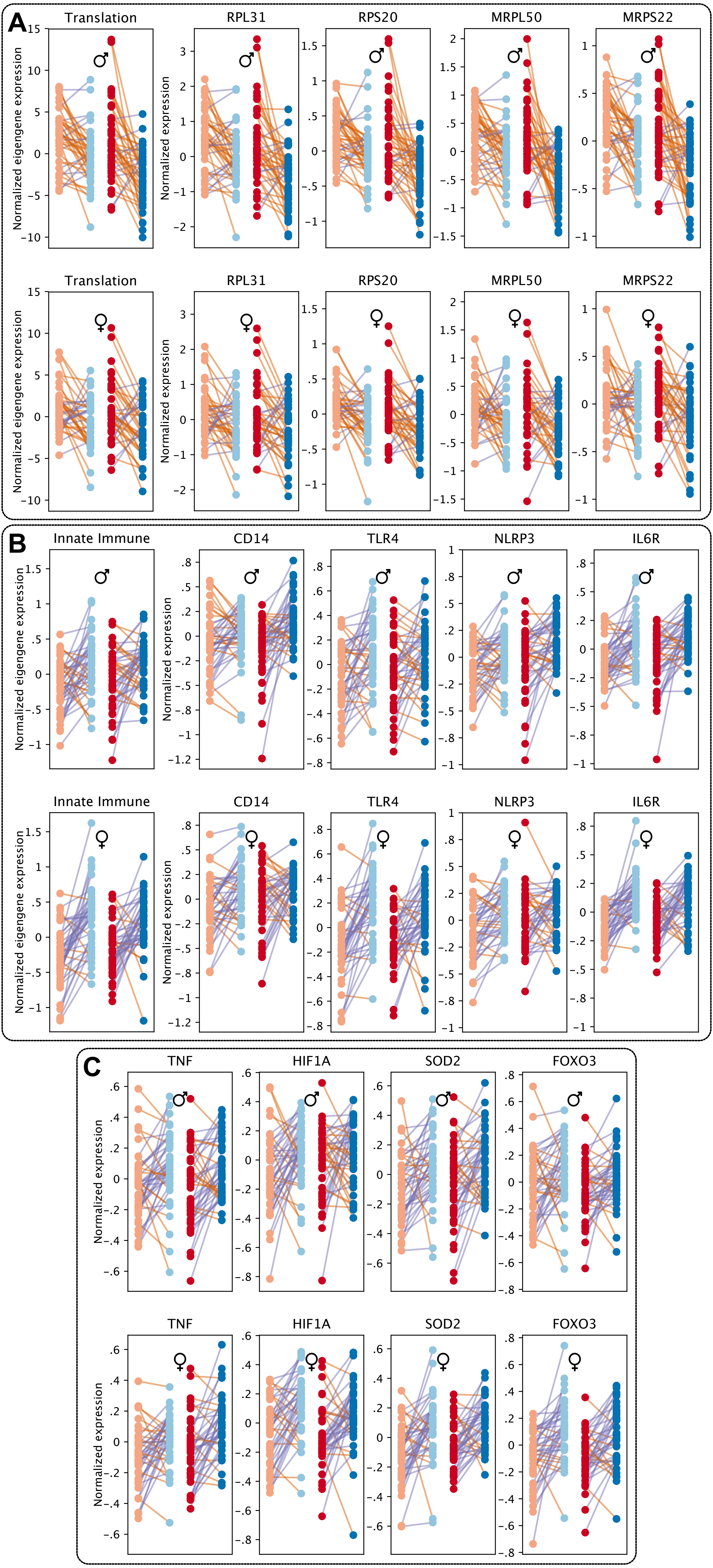

A
Translation
RPL31
RPS20
MRPL50
MRPS22
♂
♀
Normalized eigengene expression
Normalized expression
B
Innate Immune
CD14
TLR4
NLRP3
IL6R
♂
♀
Normalized eigengene expression
Normalized expression
C
TNF
HIF1A
SOD2
FOXO3
♂
♀
Normalized expression

#### SI Figure 4: Individual trajectories in PC Space

PC Trajectories of individuals in the study. In the first panel, the trajectories for individuals in response to the baseline nutrient challenge are dislplayed. Red points indicate fasted transcriptomic states, and blue points the postprandial states, projected in PC space. States from the same individual are connected with a line. The average trajectory between the two states is indicated with a thick black line. In the second panel, the same trajectories are given, but after the intervention. The third panel superimposes the baseline and post-intervention trajectories. The final panel indicates the gene expression level differences between the postprandial states (Materials and Methods).

A) For all participants
B) For males only
C) For females only

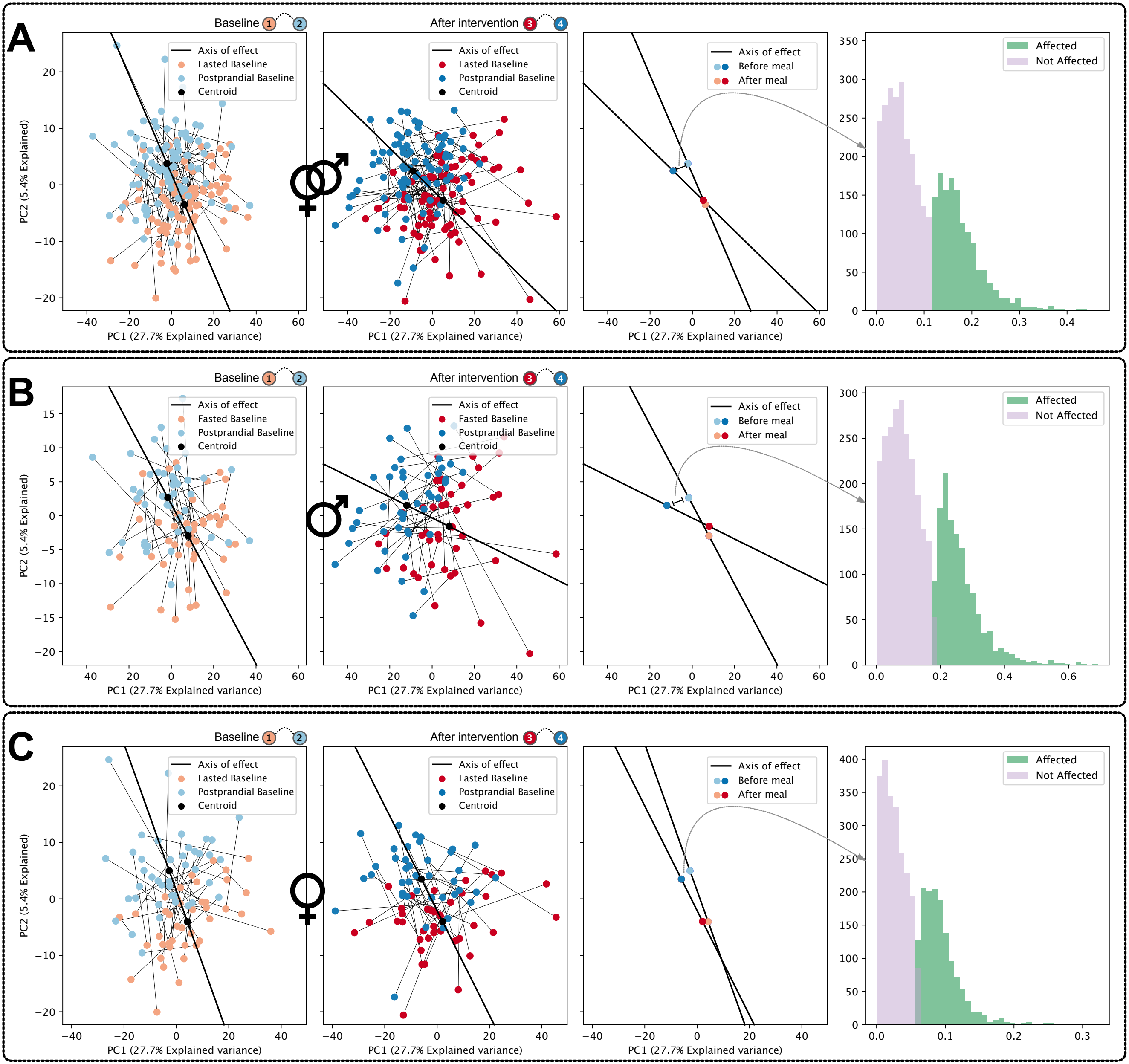

A
B
C
Baseline
After intervention
Axis of effect
Fasted Baseline
Postprandial Baseline
Centroid
Before meal
After meal
Affected
Not Affected
PC1 (27.7% Explained variance)
PC2 (5.4% Explained)

#### SI Figure 5: Change in correlation structure in males and females

For simplicity, Figure 5A showed the changes in correlation distributions for only male samples. Here we provide the same figures for males and females. The distribution of pairwise postprandial gene-response correlations at baseline (left) and post-intervention (right). The green peak represents all correlations between up-regulated genes, the blue peak between all down-regulated genes, and the brown peak between up- and down-regulated genes.

A) For males only
B) For females only

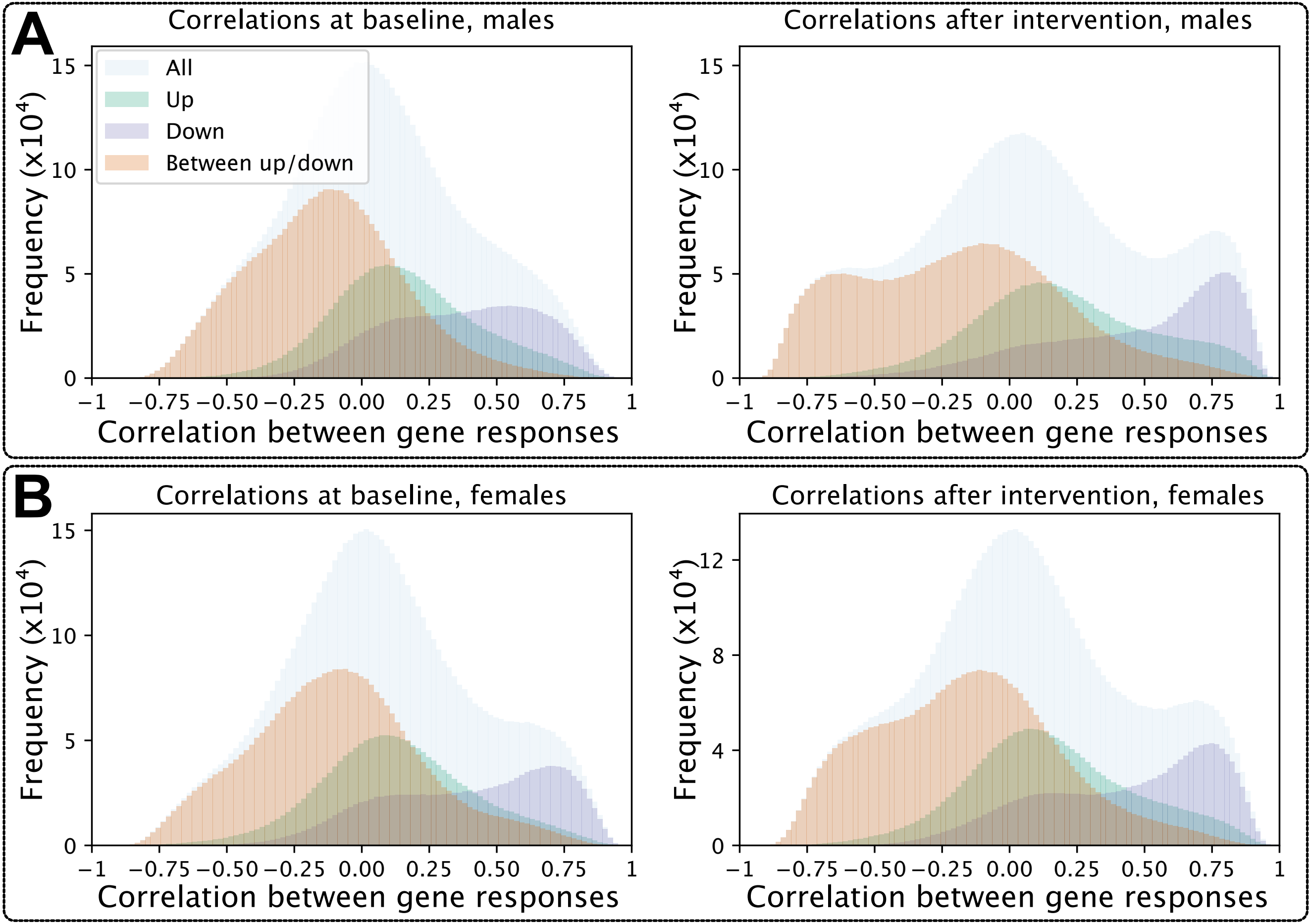
A
Correlations at baseline, males
Correlations after intervention, males
All
Up
Down
Between up/down
Frequency (x10^4)
Correlation between gene responses
B
Correlations at baseline, females
Correlations after intervention, females

#### SI Figure 6: Changes in metabolite levels by nutrient challenge

Changes in metabolite levels due to the nutrient challenge were investigated at baseline and after the intervention in men, women, and in bulk. The forest plot shows the results of the analysis described in the materials and methods.

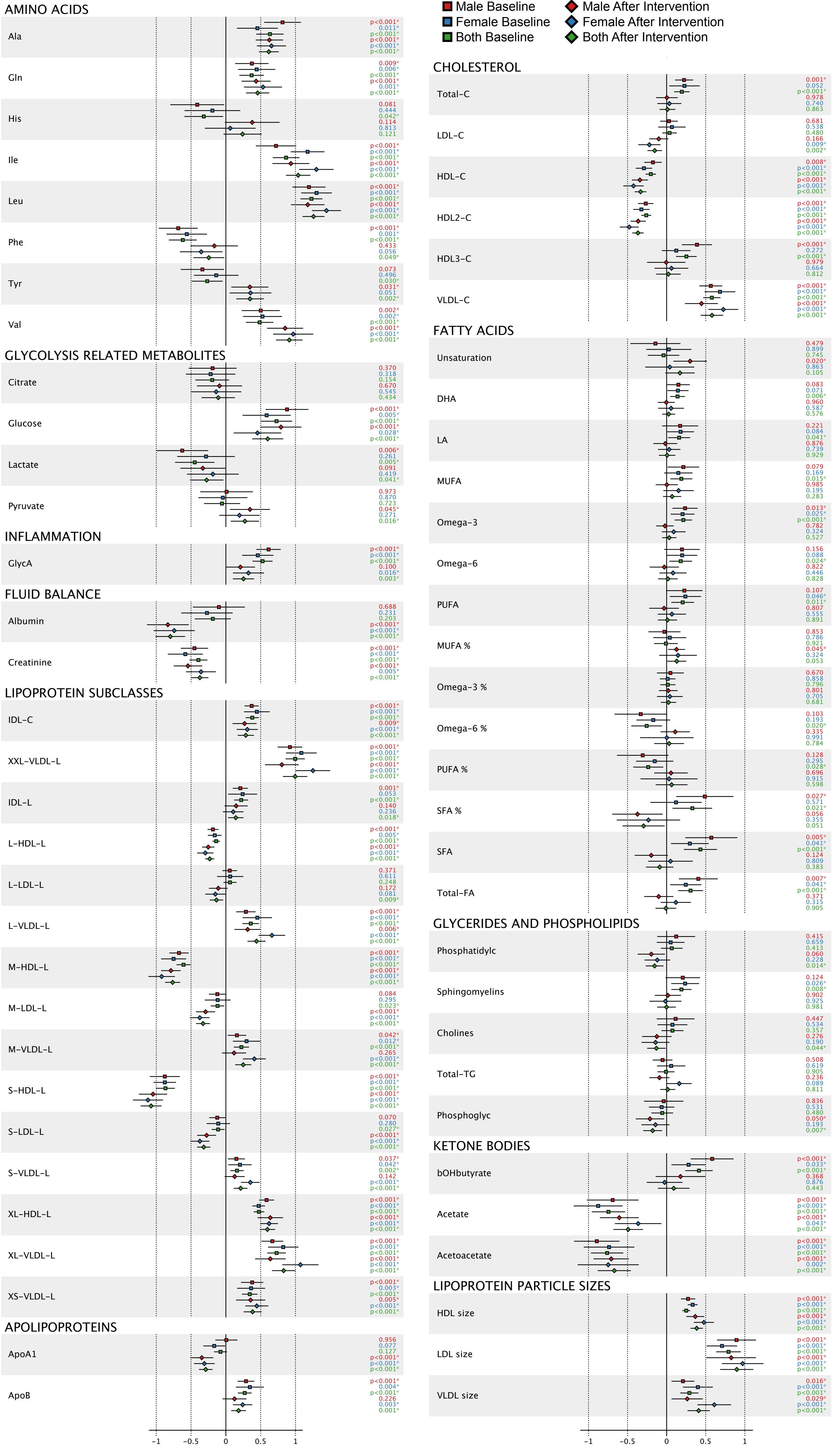

Male Baseline
Female Baseline
Both Baseline
Male After Intervention
Female After Intervention
Both After Intervention
AMINO ACIDS
Ala p<0.001* 0.011* p<0.001* p<0.001* p<0.001* p<0.001*
Gln 0.009* 0.006* p<0.001* p<0.001* 0.001* p<0.001*
His 0.081 0.444 0.042* 0.114 0.813 0.121
Ile p<0.001* p<0.001* p<0.001* p<0.001* p<0.001* p<0.001*
Leu p<0.001* p<0.001* p<0.001* p<0.001* p<0.001* p<0.001*
Phe p<0.001* 0.001* p<0.001* 0.433 0.056 0.049*
Tyr 0.073 0.496 0.030* 0.031* 0.051 0.002*
Val 0.002* 0.002* p<0.001* p<0.001* p<0.001* p<0.001*
GLYCOLYSIS RELATED METABOLITES
Citrate 0.370 0.318 0.154 0.670 0.545 0.434
Glucose p<0.001* 0.005* p<0.001* p<0.001* 0.028* p<0.001*
Lactate 0.006* 0.261 0.005* 0.091 0.419 0.041*
Pyruvate 0.973 0.870 0.723 0.045* 0.271 0.016*
INFLAMMATION
GlycA p<0.001* p<0.001* p<0.001* 0.100 0.016* 0.003*
FLUID BALANCE
Albumin 0.688 0.231 0.203 p<0.001* p<0.001* p<0.001*
Creatinine p<0.001* p<0.001* p<0.001* p<0.001* 0.005* p<0.001*
LIPOPROTEIN SUBCLASSES
IDL-C p<0.001* p<0.001* p<0.001* 0.009* p<0.001* p<0.001*
XXL-VLDL-L p<0.001* p<0.001* p<0.001* p<0.001* p<0.001* p<0.001*
IDL-L 0.001* 0.053 p<0.001* 0.140 0.236 0.018*
L-HDL-L p<0.001* 0.005* p<0.001* p<0.001* p<0.001* p<0.001*
L-LDL-L 0.371 0.611 0.248 0.172 0.081 0.009*
L-VLDL-L p<0.001* p<0.001* p<0.001* 0.006* p<0.001* p<0.001*
M-HDL-L p<0.001* p<0.001* p<0.001* p<0.001* p<0.001* p<0.001*
M-LDL-L 0.084 0.295 0.023* p<0.001* p<0.001* p<0.001*
M-VLDL-L 0.042* 0.012* p<0.001* 0.265 p<0.001* p<0.001*
S-HDL-L p<0.001* p<0.001* p<0.001* p<0.001* p<0.001* p<0.001*
S-LDL-L 0.070 0.280 0.027* p<0.001* p<0.001* p<0.001*
S-VLDL-L 0.037* 0.042* 0.002* 0.142 p<0.001* p<0.001*
XL-HDL-L p<0.001* p<0.001* p<0.001* p<0.001* p<0.001* p<0.001*
XL-VLDL-L p<0.001* p<0.001* p<0.001* p<0.001* p<0.001* p<0.001*
XS-VLDL-L p<0.001* 0.003* p<0.001* 0.005* p<0.001* p<0.001*
APOLIPOPROTEINS
ApoA1 0.956 0.077 0.127 p<0.001* p<0.001* p<0.001*
ApoB p<0.001* 0.004* p<0.001* 0.226 0.003* 0.001*
-1 -0.5 0 0.5 1
CHOLESTEROL
Total-C 0.001* 0.052 p<0.001* 0.978 0.740 0.863
LDL-C 0.681 0.538 0.480 0.166 0.009* 0.002*
HDL-C 0.008* p<0.001* p<0.001* p<0.001* p<0.001* p<0.001*
HDL2-C p<0.001* p<0.001* p<0.001* p<0.001* p<0.001* p<0.001*
HDL3-C p<0.001* 0.272 p<0.001* 0.979 0.664 0.812
VLDL-C p<0.001* p<0.001* p<0.001* p<0.001* p<0.001* p<0.001*
FATTY ACIDS
Unsaturation 0.479 0.899 0.745 0.020* 0.863 0.105
DHA 0.083 0.071 0.006* 0.960 0.587 0.576
LA 0.221 0.084 0.041* 0.876 0.739 0.929
MUFA 0.079 0.169 0.015* 0.985 0.195 0.283
Omega-3 0.013* 0.025* p<0.001* 0.782 0.324 0.527
Omega-6 0.156 0.088 0.024* 0.822 0.446 0.828
PUFA 0.107 0.046* 0.011* 0.807 0.555 0.891
MUFA % 0.853 0.786 0.921 0.045* 0.324 0.053
Omega-3 % 0.670 0.858 0.796 0.801 0.705 0.681
Omega-6 % 0.103 0.193 0.020* 0.335 0.991 0.784
PUFA % 0.128 0.295 0.028* 0.696 0.915 0.598
SFA % 0.027* 0.571 0.021* 0.056 0.355 0.051
SFA 0.005* 0.041* p<0.001* 0.124 0.809 0.383
Total-FA 0.007* 0.041* p<0.001* 0.371 0.315 0.905
GLYCERIDES AND PHOSPHOLIPIDS
Phosphatidylc 0.415 0.659 0.413 0.060 0.228 0.014*
Sphingomyelins 0.124 0.026* 0.008* 0.902 0.925 0.981
Cholines 0.447 0.534 0.357 0.276 0.190 0.044*
Total-TG 0.508 0.619 0.905 0.236 0.089 0.811
Phosphoglyc 0.836 0.531 0.480 0.050* 0.193 0.007*
KETONE BODIES
bOHbutyrate p<0.001* 0.033* p<0.001* 0.368 0.876 0.443
Acetate p<0.001* p<0.001* p<0.001* p<0.001* 0.043* p<0.001*
Acetoacetate p<0.001* p<0.001* p<0.001* p<0.001* 0.002* p<0.001*
LIPOPROTEIN PARTICLE SIZES
HDL size p<0.001* p<0.001* p<0.001* p<0.001* p<0.001* p<0.001*
LDL size p<0.001* p<0.001* p<0.001* p<0.001* p<0.001* p<0.001*
VLDL size 0.016* p<0.001* p<0.001* 0.029* p<0.001* p<0.001*
-1 -0.5 0 0.5 1

#### SI Figure 7: Modules detected in differentially expressed genes

Hierarchical clustering of differentially expressed gene responses to nutrient intake. The height of the tree refers to the 1-p correlation between genes. (i.e. a height of 0 indicates a correlation of 1, 1 indicates 0 correlation, 2 indicates -1 correlation. Different clusters are indicated in colors.

A) For females
B) For males

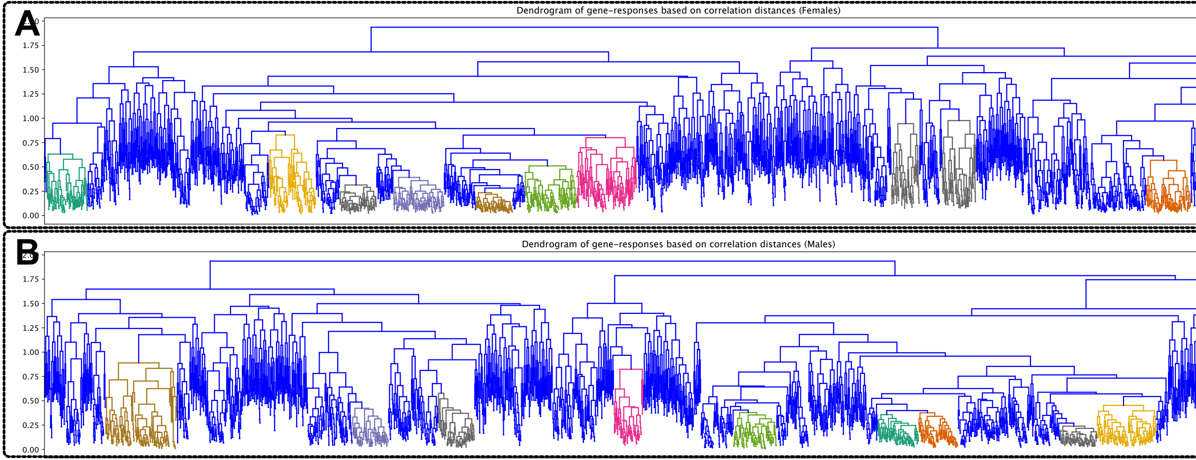
A
Dendrogram of gene-responses based on correlation distances (Females)
1.75
1.50
1.25
1.00
0.75
0.50
0.25
0.00
B
Dendrogram of gene-responses based on correlation distances (Males)
1.75
1.50
1.25
1.00
0.75
0.50
0.25
0.00
